# On Deriving Synteny Blocks by Compacting Elements

**DOI:** 10.64898/2025.12.15.694404

**Authors:** Leonard Bohnenkämper, Luca Parmigiani, Cedric Chauve, Jens Stoye

## Abstract

Genomic rearrangements are major drivers of evolution and genetic disease. However, studying rearrangements requires segmenting the genomes of interest into conserved regions, called synteny blocks, that highlight structural differences between genomes. Synteny blocks are typically defined from annotated genes or derived as a by-product of whole-genome alignments. As these procedures are heuristic and do not explicitly model rearrangements, they can obscure real variation, create false similarities, and affect phylogenetic inference.

The importance of synteny block definition has long been recognized, as shown for example by discussions on breakpoint reuse, where different definitions of synteny blocks led to different estimates of rearrangement complexity in mammalian genomes.

We present a formal framework for deriving synteny blocks directly from sequence data by partitioning genomic elements into blocks that do not contain breakpoints. A breakpoint is defined between a pair of genomes as an adjacency of shared elements that occurs in one genome but not in the other. Synteny blocks are therefore not allowed to span such boundaries, ensuring that rearrangements are not obscured. The framework is fully agnostic to the type of genomic element and applies to any genome representation expressed as sequences of elements, such as non-overlapping alignments, exact matches (MUMs/MEMs), *k*-mers, unitigs or minimizers.

We formalize two optimization problems: minimizing the total genome length after replacement by synteny blocks (the Minimum-Length Synteny Block Problem) and minimizing the number of distinct blocks (the Minimum-Size Synteny Block Problem). We show that both problems are NP-hard in general. However, when blocks are required to be collinear and to contain a shared element, we provide a linear-time algorithm with respect to the number of input elements that simultaneously minimizes both objectives. The resulting method is simple, efficient, and produces large synteny blocks without obscuring rearrangements.

## 1 Introduction

Comparative genomics aims to detect and explain similarities and differences between groups of evolutionarily related genomes; examples of such analyses include genome rearrangement studies or phylogeny reconstruction, to cite only two examples. With the availability of thousands of fully assembled genomes, from small viral genomes to large and complex genomes of eukaryotic species, it is now common for comparative studies to consider tens to hundreds of genomes at once. The size and complexity of most genomes prevents that whole-genome comparative studies can be done at the nucleotide level, and a preliminary step is to abstract the considered genomes into more compact representations over an “alphabet” of conserved genome segments called *synteny blocks*, a term coined by Pevzner and Tesler in [1].

Intuitively, synteny blocks partition genomes into non-overlapping segments, each segment belonging to a unique synteny block, such that both differences between segments from the same synteny block and similarities between segments from different synteny blocks can be ignored for the comparative analysis that is done. For example, when comparing genomes to understand their evolution in terms of large-scale genomic rearrangements, synteny blocks aim to be large homologous genome segments that are likely to have evolved from a single ancestral genome segment through nucleotide-level mutations, small-scale duplications and losses, and micro-rearrangements [1, 2, 3, 4]. In practice, synteny blocks are determined from small, highly-conserved genome segments called *elements* (e.g., orthologous genes, unitigs, local pairwise or multiple sequence alignments), that are clustered into synteny blocks according to criteria that are typically parameterized by conservation of content and order; for example, locally collinear synteny blocks would require that all genome segments in a synteny block contain the set of elements in the same order and orientation.

Since the initial comparison between the human and mouse genomes, there has been a large corpus of methods for the computation of synteny blocks, using a wide variety of approaches [5, 6, 7, 8, 9, 10, 11, 12, 13, 14, 15, 16, 17, 18, 19, 20, 21, 22, 23]. Most methods are inherently heuristic, in that they do not rely on a formal definition of what a synteny block is; we refer the reader to the recent papers [20, 21, 23] for an overview of used approaches.

As a consequence of the many ways synteny blocks can be defined, the design of algorithms to identify synteny blocks is both challenging and prone to controversy. This is well illustrated by the debate on the rate of breakpoint reuse by genomic rearrangements which stemmed from initial comparisons of the human and mouse genomes and focused on the choice of synteny blocks [24, 25, 26, 27, 28]. Additionally, in [29], which focused on rearrangement-based phylogenetic reconstruction, it was shown that “the tree inference method had less impact than the block determination method”, where impact was measured by the agreement of rearrangement-based trees with a SNP-based reference tree. In a different comparative study of synteny blocks methods [30], Ghiurcuta and Moret argued for “the need for a well founded, systematic approach to the decomposition of genomes into syntenic blocks.”

In this work, we first focus in Section 2 on introducing a formal definition of synteny blocks and on formalizing the computation of synteny blocks as two related optimization problems. We define synteny blocks as groups of genomic elements that do not contain breakpoints, where a breakpoint is defined between a pair of genomes as an adjacency of shared elements that occurs in one genome but not in the other. This definition ensures that genome rearrangements are not obscured. Our framework is fully agnostic to the type of elements and applies to any genome representation expressed as sequences of elements, such as genes, exact matches (MUMs/MEMs), *k*-mers, or unitigs. We then formalize the computation of synteny blocks as two optimization problems. The first one aims to find a set of synteny blocks minimizing the total length of the genomes after they are abstracted as (signed) sequences of synteny blocks (the Minimum-Length Synteny Block Problem), while the second one aims to find a set of synteny blocks minimizing the number of blocks (the Minimum-Size Synteny Block Problem). In Section 3, we then show that both problems are NP-hard in general, but that when blocks are required to be collinear and to contain a conserved element, i.e., an anchor present in all genome segments forming a synteny block, both problems can be solved simultaneously in linear time with respect to the input size, measured as the sum of the elements spelling all genomes. The resulting approach is conceptually simple, runs efficiently, and preserves rearrangements while producing large synteny blocks. In Section 4 we describe experiments that provide a poof-of-principle of our definition of synteny blocks and of the usefulness of our algorithms. In Section 5 we discuss the results and future research directions.

## 2 Problem Definition

We formally define *synteny blocks* and the related algorithmic problems. For the sake of exposition, we consider unichromosomal linear genomes, although our definitions naturally extend to general genomic architectures.

### Genomes and strings

Let *E* be a set of *elements*, where each element typically represents a family of genomic regions (e.g., *k*-mers, unitigs, orthologous genes).

A *string s* = *s*[1] … *s*[*n*] of length *n* over *E* is a sequence of *signed* elements, each of the form *s*[*i*] = *oe*, where *e ∈E* and *o ∈*{+,*™*}; we write sgn(*s*[*i*]) = *o* and elm(*s*[*i*]) = *e*. We use *™x* to indicate a reversal of a signed element *x*, i.e., *™*+*e* = *™e* and *™™e* = +*e*. The set of unsigned elements in a string *s* is denoted by elm(*s*) = elm(*s*[*i*]) *i* = 1, …, *n*. We denote the *substring* of string *s* between positions *i* and *j* by *s*[*i* : *j*] = *s*[*i*] … *s*[*j*].

For the moment we consider only strings in which each element occurs once, i.e., we assume that repeats have been masked. Such strings are known as *singular strings*. Our results can be generalized for arbitrary strings, with repeated elements, as outlined at the end of Section 3.2 (“Extension to Duplicates”).

### Adjacencies and Breakpoints

Rearrangements are usually characterized by their *breakpoints*. Because we aim to define blocks that reflect rearrangements and therefore breakpoints accurately, the definition of breakpoints is central to our notion of synteny blocks.

Each signed element *oe* is associated with two *extremities*: a *tail e*^*t*^ and a *head e*^*h*^. The order of these extremities depends on the sign: if *o* = +, the extremities are ordered (*e*^*t*^, *e*^*h*^); if instead *o* =*™*, the order of the extremities is reversed, (*e*^*h*^, *e*^*t*^).

The *adjacency set* of *s*, Adj(*s*), is the set of unordered pairs of extremities connecting consecutive elements:

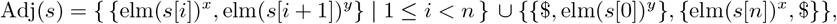

where, *x* = *h* when sgn(*s*[*i*]) = + and *x* = *t* otherwise, and *y* = *t* when sgn(*s*[*i* + 1]) = + and *y* = *h* otherwise. We call $ the *telomere*.

To compare two strings *s* and *t* with unequal content, we project them onto their common elements as follows: the *projection* of *s* onto *t*, denoted *s* _*t*_, is the string obtained from *s* by removing every *s*[*i*] for which elm(*s*[*i*]) is not contained in elm(*t*). Rearrangements can then be characterized by their *breakpoints*. The *breakpoint set* between two strings *s* and *t* is the symmetric difference between the two adjacency sets of the projections of *s* and *t* onto each other, Adj(*s*|_*t*_) and Adj(*t*|_*s*_):

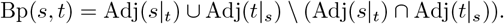

### Parsings and Partitions

We aim to define synteny blocks as a partition of the set of elements *E*. Recall that a partition *P* = {*P*_1_, …, *P*_*k*_} is a set of non-empty sets *P*_*i*_, called *parts*, such that each element is contained in exactly one part. We call the partition *P* = *{{e}* |*e ∈ E}*, the *finest partition*.

Given a partition *P*, the maximal substrings in *s* consisting of elements from the same part *P*_*i*_ form a parsing of *s*. More formally, given a string *s*, a *parsing* of *s* is a decomposition into substrings *s*[1 : *r*_1_], *s*[*r*_1_+1 : *r*_2_], …, *s*[*r*_*l−*1_ + 1 : *r*_*l*_]. Each *s*[*r*_*i−*1_ + 1 : *r*_*i*_] of a parsing is called a *phrase*. The parsing *induced* by partition *P* is the parsing, such that *s*[*i*], *s*[*i* + 1] are in the same phrase if and only if elm(*s*[*i*]) and elm(*s*[*i* + 1]) are in the same part of *P*. An example is given in Fig. 1.

**Fig. 1.**
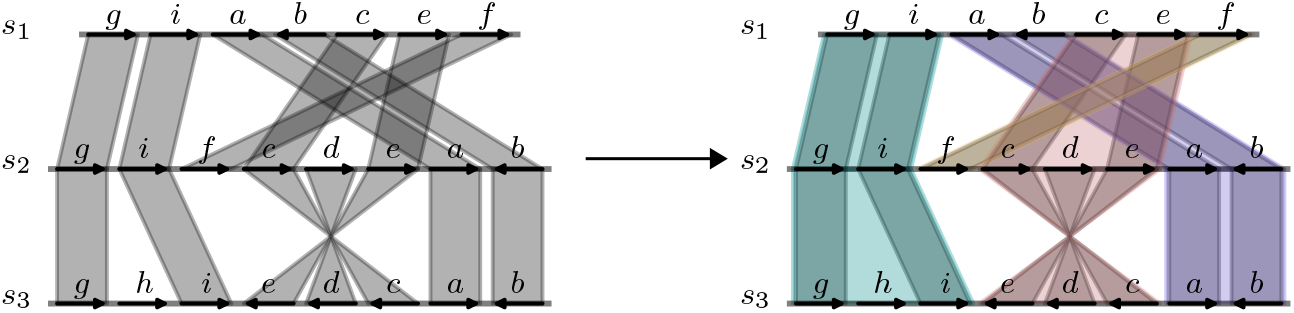
Illustration of the parsing induced by partition *P* = *{P*_1_ = *{a, b}, P*_2_ = *{c, d, e}, P*_3_ = *{f}, P*_4_ = *{g, h, i}}* for three strings; the corresponding phrases are indicated by four different colors. The parsed strings are +*g* + *i* | +*a − b* | +*c* + *e* | +*f*, +*g* + *i* | +*f* | +*c* + *d* + *e* | +*a − b*, and +*g* + *h* + *i* | *−e − d − c* | +*a − b*. One possible way of orienting the phrases is to view each as forward except *−e − d − c* in *s*_3_ which one considers backward.

In the following we define synteny blocks as partitions of *E* such that (i) phrases of distinct parts do not interleave, (ii) synteny blocks do not contain internal rearrangements (are free of breakpoints), and (iii) phrases can be assigned an orientation. The partition in Fig. 1 fulfills these properties.

### Synteny blocks

First, since our input strings are presumed to be singular, we expect synteny blocks to occur only once per string as well, uninterrupted by other blocks. An example where this is not the case is given in Fig. 2 (i). More formally: Given a part *P*_*i*_ and a signed string *s*, the *index set* of *P*_*i*_ in *s* is {*j* |elm(*s*[*j*]) *∈P*_*i*_ *}*; *P* is *absent* from *s* if its index set is empty, *present* otherwise. We call *P*_*i*_ *contiguous* in *s* if it is absent from *s* or its index set defines a single substring *s*[*i* : *j*] of *s*. We then write 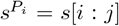 and call 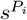 the phrase *induced* by *P*_*i*_. For a set of strings *S*, we write 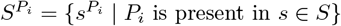.

**Fig. 2.**
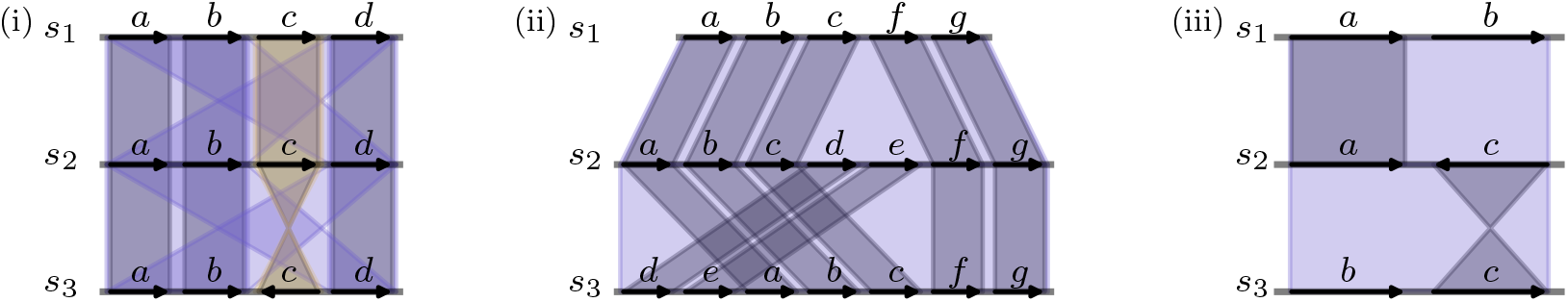
Critically undesirable properties in synteny blocks. (i) The partition {{*a, b, d}, {c}}* contains the part {*a, b, d}* which is not contiguous. (ii) This block contains a transposition of elements *d* and *e* between *s*2 and *s*3. (iii) The block {*a, b, c}* is not orientable. Trying to give this block an orientation leads to a contradiction: If it is forward in *s*1, then it is forward in *s*2 because of *a*, but also forward in *s*3 because of *b*. However, since it is forward in *s*2, it must also be backward in *s*3 because of *c*, a contradiction.

Second, since we want synteny blocks to reflect the rearrangements in the original strings, we want blocks not to contain breakpoints like shown in Fig. 2 (ii). A contiguous part *P*_*i*_ is called *breakpoint free* on string set *S* if for any two strings *s, t ∈ S*, the induced phrases 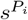 and 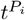 have no breakpoints with respect to each other.

Third, to reflect rearrangements in the original strings, the phrases induced by synteny blocks need to be assigned an orientation. However, this is not always possible, even when the corresponding part is breakpoint free, as shown in Fig. 2 (iii). A part *P*_*i*_ is called *orientable* on string set *S* if for each of its induced phrases in 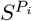, we can assign an orientation 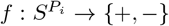, such that applying this orientation to 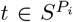, i.e., inverting it if *f* (*t*) = *−*, each element occurs only with one orientation. More formally, if 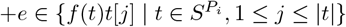, then 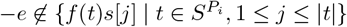 and conversely. We call *f* an *orienting function* of *P*_*i*_. For example *{a, b}* in Figure 1 is orientable. Note that negating *f*, one can always obtain another orienting function *g* of *P*_*i*_, i.e., *g*(*t*) = *−f* (*t*).

We now formally define synteny blocks and the problem(s) of finding optimal sets of synteny blocks.

#### Definition 1 (Synteny block).

*Let S be a set of strings over the set of elements E and P* = {*P*_1_, …, *P*_*k*_*} a partition of E. Part P*_*i*_ *is a* synteny block *if all of the following hold: (i) for all s ∈S, P*_*i*_ *is contiguous in s, (ii) P*_*i*_ *is breakpoint free on S, and (iii) P*_*i*_ *is orientable on S. If each P*_*i*_ *∈P is a synteny block, we call P a* synteny block partition *for S*.

For example, the partition illustrated on the right side of Fig. 1 is a synteny block partition.

As noted above, the phrases induced by blocks of *P* in each *s ∈S* define a unique parsing which we denote by *ρ*_*P*_ (*s*). We denote by *ρ*_*P*_ (*S*) the parsings of all strings in *S* induced by *P*. The parsing of *S* induced by *P* naturally defines an encoding of the strings in *S* as strings over the alphabet *{*1, …, *k}* (where *k* is the number of blocks of *P*) where, for every string *s ∈ S*, its encoding enc_*P*_ (*s*) is obtained from *s* by replacing, for every part *P*_*i*_, the phrase induced by *P*_*i*_ (if *P*_*i*_ is present in *s*) with +*i* (resp. *−i*) if *P*_*i*_ is forward (resp. reversed) in *s* according to an orienting function 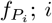 is then called the *block identifier*. For example, the encoding of the three strings in Fig. 1 is enc_*P*_ (*s*_1_) = 4 1 2 3, enc_*P*_ (*s*_2_) = 4 3 2 1 and enc_*P*_ (*s*_3_) = 4 *™*2 1.

Note that the finest partition, i.e., the one where each part contains exactly only one element, is trivially a synteny block partition. However, this is also the largest partition, both in number of parts as well as regarding the resulting encoding. Since downstream rearrangement analyses may be computationally demanding (for example, many rearrangement problems are NP-hard [31]), it makes sense to search for a smaller partition. Moreover, conceptually, a representation that requires fewer blocks is also a more parsimonious explanation for the observed rearrangements than one that requires more blocks. In each case, we are looking for the “smallest” partition. We consider two different criteria for what constitutes a partition to be the “smallest”.

*Problem 1 (Minimum-Length Synteny Blocks Problem (MLSBP))*. Given a collection of strings *S* = *{s*_1_, …, *s*_*m*_*}* over an element set *E*, find a synteny block partition *P* for *S* such that the total encoded length 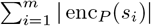 is minimized.

*Problem 2 (Minimum-Size Synteny Blocks Problem (MSSBP))*. Given a collection of strings *S* = *{s*_1_, …, *s*_*m*_*}* over an element set *E*, find a synteny block partition *P* for *S* such that the number of parts |*P* | is minimized.

Problems 1 and 2 are already meaningful in their original form, but in practice it is often desirable for the resulting synteny block partition to satisfy additional properties. We discuss these next.

### Desirable Properties of Synteny Blocks

While enforcing breakpoint-freeness and orientability is sufficient to ensure that pairwise rearrangements are not hidden within synteny blocks, it is not sufficient to do so in a global comparison on strings with unequal content. Consider Fig. 3 (i): While none of the three strings *s*_1_, *s*_2_, *s*_3_ has a breakpoint with respect to any other one, there is clearly disagreement about what the order of +*c*, +*d* and +*e* should be, indicating that there must have been a rearrangement at some point during the emergence of these “chromosomes”. Additionally, there is no multiple alignment of the phrases induced by\ this synteny block that respects all element matches. Below, we introduce the concept of *collinearity*, which ensures that such a situation does not occur by enforcing a partial ordering consistent with every phrase.

**Fig. 3.**
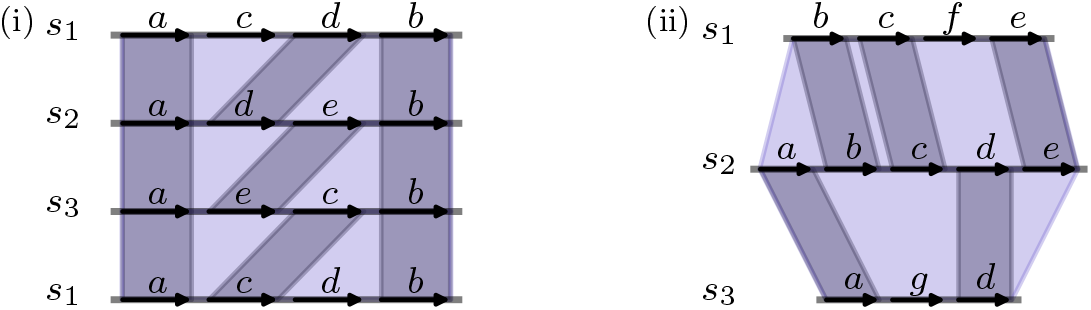
Potentially undesirable properties of synteny blocks. (i) Despite being breakpoint free, this block does not have a (partial) ordering between elements c, d and e; it is thus not collinear. (ii) The phrases induced by this block in s1 and s3 do not share any elements.

#### Definition 2 (Collinear).

*Let S be a set of strings. A synteny block P*_*i*_ *with orienting function f is* collinear *in S if there exists a partial order ≺ on the signed elements in P*_*i*_ *such that, for every s ∈ S:*

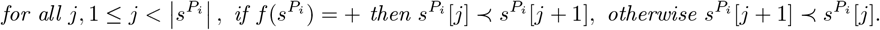

Collinearity implies breakpoint-freeness, as stated in the following lemma, proved in Appendix A.

#### Lemma 1.

*Let S be a set of strings over element set E, and P*_*i*_ *a contiguous, orientable part of a partition on E. If P*_*i*_ *is collinear in S, then P*_*i*_ *is breakpoint-free on S*.

Additionally, one may wish to avoid cases where completely unrelated phrases are induced by the same block, such as in Fig. 3 (ii). This can create false signals of homology or rearrangement. Other problems, such as inconsistent orientation (Fig. 2 (iii)) can also arise when phrases induced by a block share no common elements. To prevent this, we require an *anchor* element.

#### Definition 3 (Anchored).

*Let S be a set of strings and P* = *{P*_1_, …, *P*_*k*_*} be a set of synteny blocks for S. Recall that* 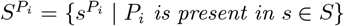. *We say that P*_*i*_ *is* anchored *if* 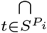 elm(*t*) *is not empty. We call any element in* 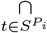 elm(*t*) *an* anchor *of P*_*i*_. *We say that P is anchored if all synteny blocks P*_*i*_ *are anchored*.

For readers familiar with the concept of “core elements” in pangenomics, we clarify that the concept of an anchor is distinct from the concept of a core element. An anchor does not need to appear in each input string, but only in those input strings where the respective block is present. In principle, even an element that occurs only in a single genome of a larger set of genomes, may be an anchor, whereas such an element is typically not considered part of the core.

Requiring anchored synteny blocks has additional benefits, some of which will be discussed later. Importantly for subsequent analyses, it enables a bijection between breakpoints in the input strings and in the encoded strings which implies preservation of genome rearrangement distances.

#### Theorem 1.

*Let P be an anchored synteny block partition for a set S of singular strings. For each pair of strings s, t ∈ S there is a bijection β*_*s,t*_ *between extremities in* Bp(*s, t*) *and in* Bp(enc_*P*_ (*s*), enc_*P*_ (*t*)), *such that*

- *for each breakpoint {a, b} ∈* Bp(enc_*P*_ (*s*), enc_*P*_ (*t*)), *{β*_*s,t*_(*a*), *β*_*s,t*_(*b*)*} is a breakpoint of s, t*,
- *for each breakpoint* 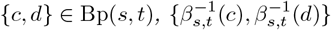 *is a breakpoint of* enc_*P*_ (*s*), enc_*P*_ (*t*).

We prove this theorem in Appendix B. The bijection between breakpoints is a powerful property, as many rearrangement models – such as the Reversal [32], Hannenhalli-Pevzner [33], Single-Cut-or-Join [34] and Double-Cut-and-Join [35] models – consider only breakpoints, not conserved adjacencies. This means that distances under such models are preserved for an anchored synteny block partition.

Finally, we propose the following variants of Problems 1 and 2, restricted to synteny blocks with these desirable properties, which play a central role in our results.

*Problem 3 (Collinear anchored-MSSBP)*. Given a collection of strings *S* = {*s*_1_, …, *s*_*m*_} over an element set *E*, find a synteny block partition *P* for *S* in which all blocks are collinear and anchored, and which is optimal with respect to the partition size, as defined in Problem 1.

*Problem 4 (Collinear anchored-MLSBP)*. Given a collection of strings *S* = {*s*_1_, …, *s*_*m*_} over an element set *E*, find a synteny block partition *P* for *S* in which all blocks are collinear and anchored, and which is optimal with respect to the encoded length, as defined in Problem 2.

## 3 Methods

With these definitions in place, we now consider the computational complexity of the problem and propose an optimal algorithm for the restricted version. First, we prove that both Problem 1 (MLSBP) and Problem 2 (MSSBP) are NP-hard in general. Then we consider Problems 3 and 4 where blocks must be collinear and anchored. Under these constraints, both problems have the same optimal solution, and this solution can be computed in linear time by a simple greedy merging procedure.

### 3.1 Finding a Smallest Synteny Block Partition is NP-hard

The optimization Problems 1 and 2 admit no polynomial time solution unless P = NP. We establish this via two polynomial-time reductions: MLSBP is shown NP-hard by a reduction from Vertex Cover, and MSSBP is shown NP-hard by a reduction from SAT. The detailed proofs appear as Theorems 5 and 6 in Appendices C and D, respectively, from which the following result follows.

#### Theorem 2.

*Problems 1 and 2 are NP-hard*.

### 3.2 Finding a Smallest Collinear, Anchored Synteny Block Partition is Polynomial

Although the unrestricted problems are hard, when adding the constraints that blocks should be collinear and have (at least) one anchor, the problem is much easier to solve. In fact, as we show in this section, the solution for both problems is the same.

We first formalize the notion that when every occurrence of an element forms an adjacency with the same element, they should belong to the same synteny block.

#### Definition 4 (Unique neighbor).

*We say that extremity a*^*x*^ *has a* unique neighbor *extremity b*^*y*^ *on string set S, for some elements a, b and some x, y ∈ {h, t}, if {a*^*x*^, *b*^*y*^*} is the only adjacency involving a*^*x*^ *on S. We write a*^*x*^ *→ b*^*y*^. *When the concrete extremities x, y are irrelevant, we simply write a → b*.

*For two parts P*_*i*_, *P*_*j*_ *of partition P, we write P*_*i*_ *→ P*_*j*_ *if in any encoding* enc_*P*_ (*S*), *i → j*.

Note that the unique neighbor relationship is directed, i.e., for *a*→ *b*, whenever *a* occurs, *b* occurs adjacent to it, but the converse is not necessarily true. This is in contrast to the concept of compaction employed in compacted de Bruijn graphs, where two *k*-mers *a, b* are placed in the same unitig if *a* always occurs adjacent to *b*, but also *b* always occurs adjacent to *a*. In our notation, this would be *a*→ *b and b* →*a*. We also note that if *i*^*x*^ →*j*^*y*^ for some *x, y ∈*{*h, t}* in an encoding of a partition *P* with *P*_*i*_, *P*_*j*_ *∈P*, we can always choose the orienting functions of *P*_*i*_, *P*_*j*_, such that *i*^*h*^ →*j*^*t*^.

We now show that one can create a set of collinear anchored synteny blocks by iteratively merging smaller (also collinear anchored) synteny blocks, where one of them is a unique neighbor of the other.

#### Lemma 2.

*Let P*_*i*_, *P*_*j*_ *be collinear synteny blocks of partition P with anchors c*_*i*_, *c*_*j*_, *such that P*_*i*_ *→ P*_*j*_. *Then P*_*i*_ *∪ P*_*j*_ *is a collinear synteny block with anchor c*_*j*_.

*Proof*. In the encoding enc_*P*_ (*s*), *P*_*i*_, *P*_*j*_ are represented by *i, j*. W.l.o.g. let *i*^*h*^ →*j*^*t*^. Given that *P*_*i*_, *P*_*j*_ are synteny blocks, the only strings relevant to whether *P*_*i*_ *∪P*_*j*_ is a synteny block are those where both *P*_*i*_ and *P*_*j*_ are present. Since *i*^*h*^ →*j*^*t*^, in all strings where *P*_*i*_ and *P*_*j*_ are present, they occur in the same orientation and their induced phrases are adjacent. Thus *P*_*i*_ *∪P*_*j*_ is a synteny block and concatenating the partial orders of *P*_*i*_ and *P*_*j*_, such that *e ≺d* for all *e ∈P*_*i*_, *d ∈P*_*j*_ is a partial order that satisfies Definition 2 for each phrase. Moreover, in each string where *P*_*i*_ is present, *P*_*j*_ is also present, therefore anchor *c*_*j*_ is also present. Thus, *P*_*i*_ *∪ P*_*j*_ has anchor *c*_*j*_.

This strategy can then be used to construct any collinear anchored synteny block partition.

#### Theorem 3.

*Any collinear anchored synteny block partition P* = *{P*_1_, …, *P*_*k*_*} on element set E* = *{e*_1_, … *e*_*l*_*} can be constructed by iteratively merging pairs of parts Q∪ R, where Q → R, starting from the finest partition {{e*_1_*}*, …, *{e*_*l*_*}}*.

We provide here a high-level sketch of a constructive proof, with full details in Appendix E. Recall that an anchored collinear block must have an anchor *c* and a partial order *≺* which describes the order of elements in each phrase. Starting from the anchor *c*, regard the closest element *e* to *c* in the partial order, such that no other element *d* exists with *c≺ d ≺e*. Then in each string where *e* occurs, *c* must also occur (since *c* is an anchor) and moreover, because there can’t be any element *d* between *c* and *e, c* is the unique neighbor of *e*. This argument can be iterated using a topological sorting of the block elements. One can thus construct any collinear anchored synteny block by merging elements with their unique neighbors.

To characterize optimal sets of synteny blocks, it is helpful to keep track of only one anchor in case there are multiple anchors per block.

#### Definition 5 (Canonical anchor).

*For a collinear anchored partition P* = {*P*_1_, …, *P*_*k*_} *on strings S on element set E, we call the smallest anchor c*_*i*_ *∈P*_*i*_ *according to an (arbitrary) total order on E the* canonical anchor *of P*_*i*_.

The canonical anchors identify the parts that they belong to, since each phrase must contain them. It is therefore possible to find an encoding of the input string set by simply removing all elements that are not canonical anchors.

#### Observation 1

*Let P* = {*P*_1_, …, *P*_*k*_*} be a collinear anchored synteny block partition of element set E with canonical anchors c*_1_, … *c*_*k*_ *on string set S. Then the string set S*^*′*^ *where each element but c*_1_, …, *c*_*k*_ *is removed and then each* +*c*_*i*_ *(−c*_*i*_*) is replaced by* +*i (−i) is an encoding for P*.

We can then construct a partition *P* where no further parts can be merged by iteratively unifying synteny blocks. The corresponding encoding *S*^*∗*^ can then be created by removing elements that are not canonical anchors. To that end, we denote by *S\e* for a string set *S* the operation that removes each occurrence of element *e* in *S*. Pseudocode is given in Algorithm 1 and a visual representation in Appendix G, Fig. 8.

#### Algorithm 1

Generate a partition *P* of collinear, anchored synteny blocks and its encoding *S*^*∗*^.

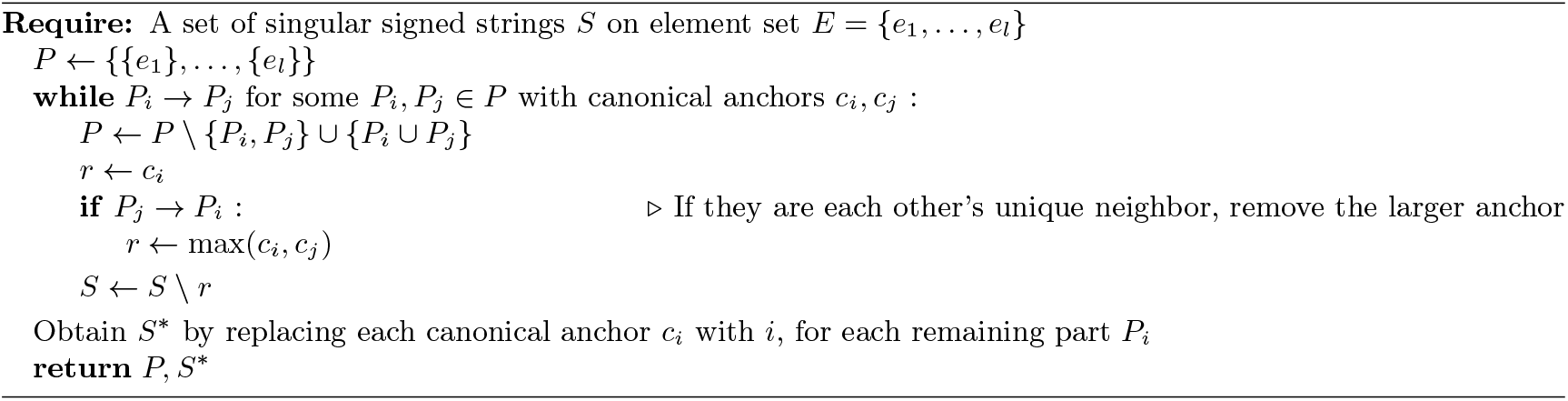

The way elements are merged here is similar to other compaction algorithms, such as the compaction step in maf2synteny [17]. However, while these algorithms are heuristics, Algorithm 1 is optimal for collinear, anchored MLSBP and MSSBP, as we show in the following. We begin by showing that, in the first iteration of the while loop, the removed element *r* from *S* cannot be a canonical anchor.

#### Lemma 3.

*Consider two elements e, d with e* →*d. If d ↛ e or d < e and e is a canonical anchor in an anchored, collinear partition P* = {*P*_1_, …, *P*_*k*_ *}, then there is a shorter partition P* ^*′*^ *with a shorter encoding than P, such that P* ^*′*^ *is also anchored and collinear*.

*Proof*. (By contradiction) Assume that *e* is a canonical anchor of part *P*_*i*_. Note that because *e → d* and *d < e or d ↛ e, e* cannot be the canonical anchor of a block which contains *d*, i.e. *d ∉P*_*i*_. Let *d ∈ P*_*j*_ with canonical anchor *c*. Then *P*_*i*_ *→ P*_*j*_ (because *e → d*), that is *P*_*i*_ *∪ P*_*j*_ is a collinear synteny block and has anchor *c* according to Lemma 2. Observe that *P* ^*′*^ = *P\{P*_*i*_, *P*_*j*_*} ∪ {P*_*i*_ *∪ P*_*j*_*}* has both fewer parts and a shorter encoding than *P*.

Note that removing an element also does not change whether the phrases induced by a block are contiguous or collinear, and as long as one anchor remains, the block also remains anchored. Thus, any optimal solution *P* on *S* still has an equivalent solution on *S\r*.

#### Observation 2

*Given a set of strings S* = *{s*_1_, …, *s*_*m*_*} and a collinear anchored partition P* = *{P*_1_, …, *P*_*k*_*}, let P \ e be the set of parts and S \ e be the set of strings obtained by removing an element e that is not a canonical anchor. Then P \ e is a collinear anchored synteny block partition on S \ e*.

Let *S*_*q*_ be *S* in the *q*-th iteration of Algorithm 1. It follows from Observation 2 and Lemma 3 that any optimal solution on *S*_*q*_ still has a (possibly suboptimal) equivalent solution on *S*_*q*+1_ and the element *r* removed in *S*_*q*+1_ never is a canonical anchor of an optimal solution on *S*_*q*_. Then any optimal solution on *S*^*∗*^ is at most as large as any optimal solution on *S*.

#### Corollary 1.

*Let S*^*∗*^ *be the string set produced by Algorithm 1*.

*Let* 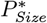, *P*_*Size*_ *be the collinear, anchored partitions on S*^*∗*^ *and on Srespectively, such that* 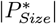 *and* |*P*_*Size*_| *are minimized. Then* 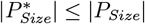.

*Let* 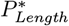, *P*_*Length*_ *be collinear, anchored partitions on S*^*∗*^ *and on S respectively, such that the respective encoding lengths* 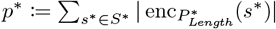 *and* 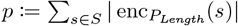 *are minimized. Then p*^*∗*^ *≤ p*.

Note that on *S*^*∗*^ there are no further elements with unique neighbors. On such a string set, Theorem 3 implies that the only collinear anchored synteny block partition on *S*^*∗*^ is the finest partition. Since |*P* | is the number of elements in *S*^*∗*^ and 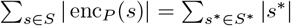, the partition constructed by the algorithm is optimal for both MLSBP and MSSBP.

#### Theorem 4.

*Algorithm 1 constructs a partition P that is a solution to Problems 3 and 4*.

This shows that in the restricted case Problems 3 and 4 are in fact equivalent. However, given the differences in the hardness proofs, we conjecture that in general Problems 1 and 2 are distinct.

With the correctness of the greedy algorithm established, we analyze its computational complexity. Let *L* denote the total number of extremity occurrences across all genomes, counted with multiplicity.

#### Lemma 4.

*Algorithm 1 admits an implementation which is linear in time and space with respect to the input genome size L*.

A detailed proof is provided in the Appendix F; in the following we give a brief summary. Algorithm 1 repeatedly merges elements that have a unique neighbor until no further merges are possible. Initially, all extremities with a unique neighbor are added to a queue. While the queue is not empty, the algorithm removes one extremity *a*^*x*^, determines its unique neighbor *b*^*y*^ (with *a* → *b*), and merges the corresponding element *a* into *b*. All extremities *c*^*z*^ that were adjacent to the opposite extremity of *a*^*x*^ are reconnected to *b*^*y*^. If this update causes any of these neighboring extremities to have a unique neighbor, they are added to the queue. The algorithm terminates when the queue is empty. The complexity analysis has three main components: the number of iterations of the main while loop, the cost of updating adjacencies following a merge, and the cost of detecting new unique neighbors. We show that the while loop considers each adjacency at most twice, each corresponding to a possible merge between two elements. Merging an element *a* into *b* affects only the occurrences of *a*, and since merged elements are never reinserted, the total update cost remains linear in *L*. Finally, checking for new unique neighbors can be done in amortized linear time with respect to *L*, as each adjacency list is scanned only until one or two non-merged elements are found.

### Extension to Duplicates

We propose two extensions to handle duplicates. First, we introduce an approach that marks duplicated elements and prevents them from merging with any other element. In this way, breakpoints are fully preserved (by Theorem 1). We refer to this variant as the “breakpoint (BP) bijection” mode. Second, to allow for smaller block representations, we permit merging with duplicated elements. The theoretical implications are explored in Appendix G. However, it is not possible to define breakpoints between duplicate elements without disambiguating copies of duplicates, which in turn depends on which rearrangement model is used and is typically NP-hard [36, 37]. We therefore require in the duplicate theory presented in Appendix G that phrases induced by blocks contain each element at most once, i.e., each phrase is a singular string of elements. This allows to define a block as breakpoint-free if none of its phrases have breakpoints with respect to each other. However, this does not guarantee that global breakpoints are preserved, because Theorem 1 cannot be defined for duplicates without a concrete rearrangement model. And even singular elements with a breakpoint between them can appear in the same block if they are merged with duplicates as we show in Appendix Fig. 7. Nonetheless, this theory guarantees that the phrases induced by a block are breakpoint free with respect to each other, which implies in particular that any non-duplicate block does not obscure any rearrangements. We refer to this mode in the following as the “duplicates” mode.

These two approaches mirror the concepts of *global collinearity* and *local collinearity*, found in the existing literature on synteny blocks [20]; the “breakpoint (BP) bijection” mode preserves breakpoints globally, while the “duplicates” mode preserves breakpoints locally.

## 4 Results

We implemented the linear version of Algorithm 1 and made it available at github.com/gi-bielefeld/mice. We refer to this implementation as *Markers Inferred by Compacting Elements (MICE)*. We tested three distinct modes of the algorithm. Since the theory presented here is based on singular elements, the default mode (“MICE”) ignores duplicates as elements, but may still create blocks that span positions with duplicate elements. As described in the previous section, we offer two modes with different guarantees for duplicates. “MICE duplicates” filters out elements which occur more than 10 times per genome while allowing merges with duplicates. “MICE BP bijection” on the other hand does not permit merging with or over any duplicate element.

For our proof-of-concept experiments, we conceptually use 31-mers as elements, since *k*-mers are a well-known annotation-free and frequently used type of object in genomics and *k* = 31 is among the most frequent settings of this parameter. Since our compaction is a generalized version of unitigs, we used unitigs directly as input, not individual *k*-mers. We generated unitigs using ggcat [38]. This value for *k* is slightly higher than for SibeliaZ, which we describe next. However, for MICE the value of *k* primarily defines the threshold above which rearrangements are considered, not sensitivity for similarity. We benchmarked MICE against state-of-the-art synteny block detection methods, described in the following.

*SibeliaZ* [20] is a heuristic algorithm that also works on elements (usually *k*-mers), selecting a block if there is a strong enough “backbone” of *k*-mers. It is currently one of the fastest tools to generate synteny blocks. In each case, we used the recommended *k*-value (*k* = 15 for bacteria, *k* = 25 otherwise) and two different *k*-mer frequency thresholds *a*, namely *a*_low_ = 2 *×m* and *a*_high_ = 2 *×*10*× m*. According to its documentation, SibeliaZ with *a*_low_ should ignore duplicates, thus behaving like the default mode of MICE. SibeliaZ with *a*_high_ should be able to recover duplicates that occur up to 10 times and therefore behave more similarly to MICE duplicates or MICE BP bijection.

*Minigraph-Cactus* [39] is a workflow that builds on minigraph [40] using the Cactus framework [41, 42], which is an alignment-based, optimization-free way to construct pangenome graphs and synteny blocks.

### Datasets

We used five pangenomic datasets to assess the performance of our method, namely a selection of 56 *Y. pestis* assemblies, 48 *E. coli* assemblies, all 29 currently available *S. cerevisiae* assemblies with NCBI assembly level “complete”, all 5 currently available *A. thaliana* assemblies with NCBI assembly level “complete” as well as the 16 *M. musculus* genomes studied in the initial publication of SibeliaZ [20]. We give details to these datasets in Appendix H, Table 1.

### 4.1 Benchmark

#### Runtime

We benchmarked MICE against SibeliaZ. For both methods we only evaluated the synteny block generation step, that is, we measured the time it takes to convert elements as input to blocks as output. To that end, we provided MICE with a GFF-format file representing unitigs in the ggcat graph and we provided SibeliaZ with the binary TwoPaCo graph and skipped SibeliaZ’s alignment step. We did not include Minigraph-Cactus in this evaluation as the costly alignment step is central to the Minigraph-Cactus workflow. The runtimes for Minigraph-Cactus in each case were slower (see also Fig. 9 in Appendix H), but are not directly comparable to the measurements reported here.

We ran all algorithms on a single thread on a cluster machine. The results are visualized in Fig. 4. We see that despite being an exact algorithm with strict theoretical guarantees, MICE achieves similar performance to SibeliaZ, even outperforming the version with a high abundance filter. However, for both tools we observed that a significant portion of the runtime was spent on loading the input data from disk. We therefore see major potential for performance improvement by integrating ggcat directly with MICE. Because not all SibeliaZ runs finished within 24 hours on the *M. musculus* dataset, in the following experiments we only used the first 4 of the 16 assemblies of this dataset.

**Fig. 4.**
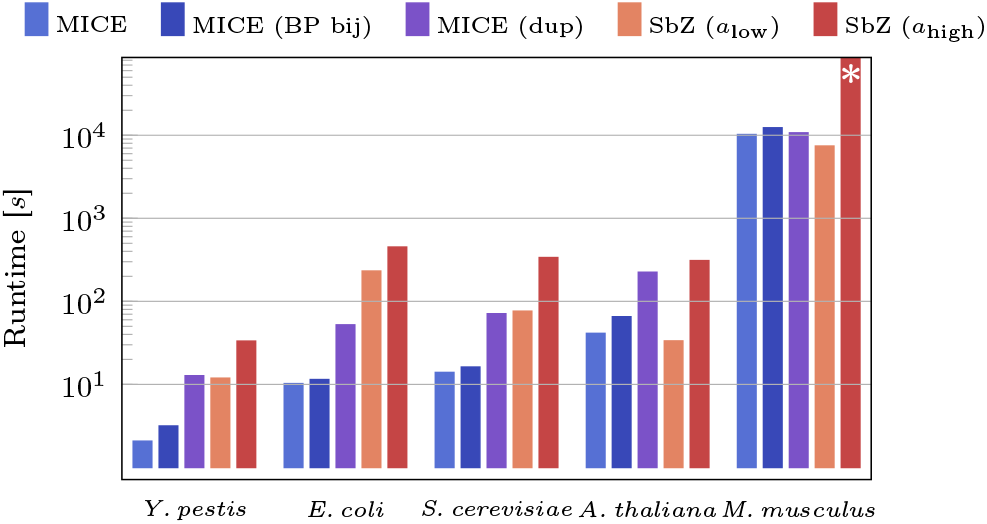
Runtimes of three MICE variants (MICE, breakpoint-bijection mode “MICE (BP bij)”, and duplicate mode “MICE (dup)”) and two SibeliaZ variants (SbZ with *a*_low_ and *a*_high_) on five benchmark datasets, measured with `/usr/bin/time` wall-clock time. The asterisk in SbZ (*a*_high_) for *M. musculus* indicates that it exceeded the 24-hour limit.

#### Block contiguity

We proceeded to evaluate the coverage and contiguity of SibeliaZ, Minigraph-Cactus and MICE on the benchmark datasets. By default, SibeliaZ restricts its output to blocks of 50 basepairs or longer. To ensure a fair comparison, we therefore included only blocks of at least 50 basepairs for all tools in this analysis. We examined how many blocks it takes to cover a certain fraction of the input genomes. As a summary, we give the coverage and encoding sizes in Table 2 in Appendix H and the respective N50, N75 and N90 values in Fig. 5. Here N*x* is defined as the length threshold with which one can cover *x*% of the total genome lengths using only sequences of length N*x* or higher. We show details in Fig. 10 of Appendix H. We see that for all datasets, the default version of MICE covers the same or more positions with fewer and larger blocks than SibeliaZ and Minigraph-Cactus. Running in duplicate and BP bijection mode produces slightly shorter blocks on low duplicate datasets, but has problems covering genomes with large numbers of duplicates.

**Fig. 5.**
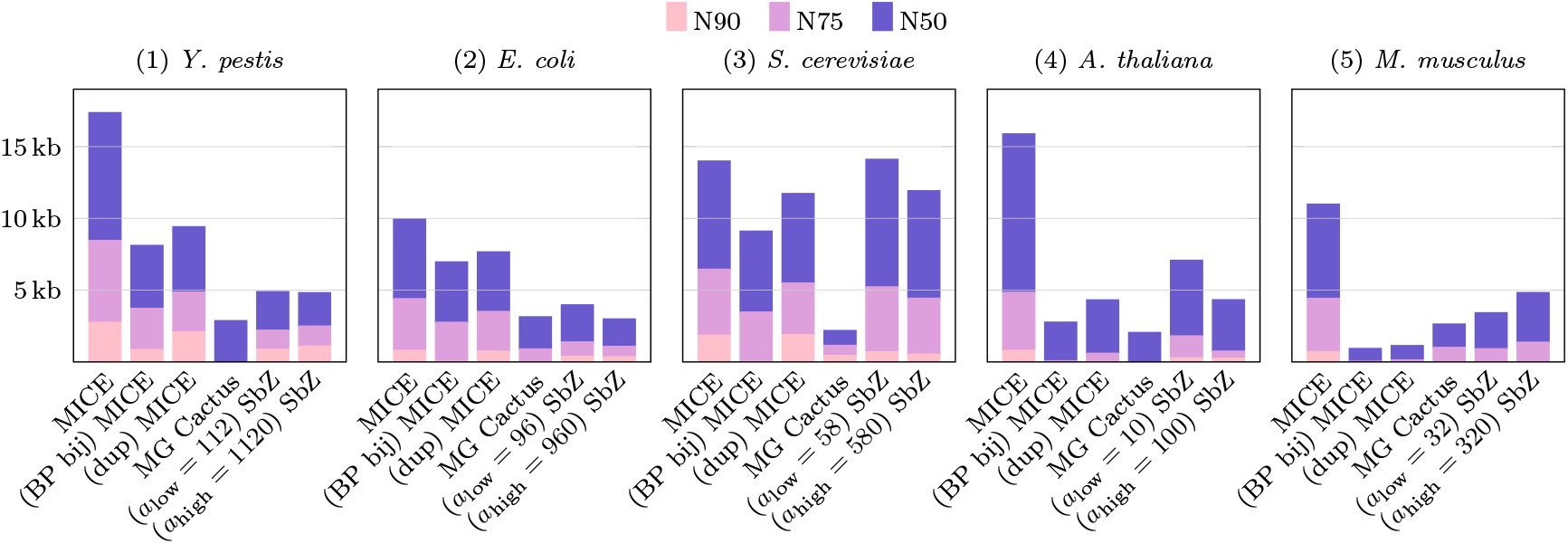
N50, N75 and N90 values for blocks generated by the tools (on the *x*-axis): MICE, MICE in breakpoint bijection mode (“(BP bij) MICE”), MICE with duplicates (“(dup) MICE”), Minigraph-Cactus (MG Cactus) and SibeliaZ (SbZ), evaluated on five benchmark datasets.

Nonetheless, the coverage improvement of the default mode in these genomes is even more noticeable. Additionally, examining the proportion of shared blocks between pairs of genomes (Appendix H, Fig. 11) shows that the high coverage of MICE does not come at the expense of significantly reducing sensitivity in finding shared regions.

### 4.2 Rearrangement Detection

While it is desirable for synteny blocks to be large and have high coverage, they must also avoid obscuring true rearrangements or create new ones. To assess this, we measured precision and recall for each method and for four of the five datasets. We excluded the *M. musculus* dataset because determining breakpoints on the original elements in this dataset was too computationally expensive.

Breakpoints involving duplicated elements are not well-defined without assuming a specific rearrangement and matching model. Therefore, for this analysis, we discarded any element that appeared as a duplicate in at least one genome of the dataset. For each dataset, we removed duplicated elements, compared all pairs of genome sequences, projected their sequences onto their shared 31-mers, and computed the set of breakpoints. We refer to this set as the element breakpoints.

We define a breakpoint as unrecoverable if an element breakpoint falls entirely within a block, making it undetectable without inspecting the elements inside the block. For each block produced by a tool, we identified the elements whose coordinates lie within the block’s coordinate range. For every such block, we examined all pairs of contained elements (*a, b*). If (*a, b*) constitutes an element breakpoint but the block merges them, this breakpoint is counted as a false negative. In Appendix I, we compare this definition to a more relaxed variant in which all elements assigned the same block identifier are unified; the resulting numbers of unrecoverable breakpoints are very similar.

The set of true positives consists of all element breakpoints that are not obscured by any block. Conversely, false positives are defined as breakpoints between two blocks *x* and *y* that do not correspond to a breakpoint in the element set. To identify these, we again compare each pair of genomes and project their sequences onto the shared blocks. A breakpoint between *x* and *y* is counted as a false positive if there is no supporting element breakpoint between any element in block *x* and any element in block *y*.

Using these definitions, recall is computed as the fraction of element breakpoints that remain detectable after block construction, i.e., the proportion of true positives among all element breakpoints. Precision is computed as the fraction of true positives breakpoints among all the one recalled, i.e., the proportion of true positives among all predicted breakpoints. Figure 6 reports precision and recall for all evaluated methods.

**Fig. 6.**
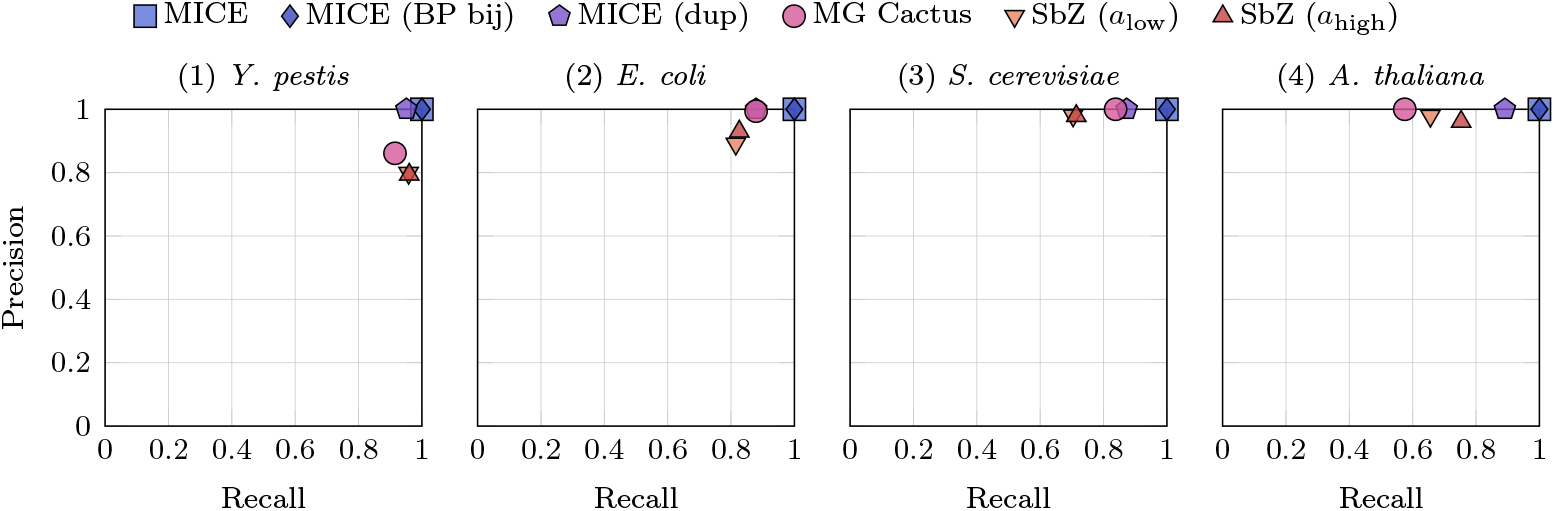
Precision and recall for all methods across four datasets. A false negative is a pair of elements that forms a breakpoint in a pair of genomes but is placed within the same block, thereby obscuring the breakpoint. True positives are element breakpoints that remain detectable after block construction. False positives are breakpoints implied by block pairs that have no supporting element breakpoint between the corresponding elements.

By construction, MICE and MICE (BP bij) achieve 100% precision and 100% recall (Theorem 1). MICE (dup) also attains 100% precision by construction. Its recall is the highest among the remaining methods on all datasets except *Y. pestis*, where it is comparable to the other tools, though still below the recall of MICE and MICE (BP bij). MG Cactus achieves the highest precision among the non-MICE methods, reaching 100% on *S. cerevisiae* and *A. thaliana*. Its recall ranges between 84% and 92% on all datasets except *A. thaliana*, where it decreases to 58%. SibeliaZ (*a*_low_) and SibeliaZ (*a*_high_) show similar performance. Both achieve precision between 80% and 98% and recall between 70% and 96%. As with MG Cactus, recall drops on *A. thaliana*, where it decreases to 66%. Overall, SibeliaZ (*a*_low_) has marginally higher precision and marginally lower recall than SibeliaZ (*a*_high_).

Additionally, to illustrate how breakpoints can become unrecoverable, we selected three *E. coli* genomes for a small-scale visualization. We applied all five methods and chose a representative locus to compare their behavior. This analysis is presented in Appendix J.

## 5 Discussion

We introduced a formal framework for deriving synteny blocks by compacting genomic elements in a way that preserves rearrangements. To our knowledge, this is the first such framework. We defined the MSSBP and MLSBP problems and showed that both are NP-hard in general. Yet, when blocks are required to be collinear and contain an anchor element, both objectives can be optimized simultaneously by a linear-time algorithm. Our experiments indicate that, using 31-mers as elements, this algorithm is competitive with state-of-the-art synteny block construction methods while preserving all rearrangements of unique elements present in the input data. Nonetheless, which elements to use as input for different research objectives, is an open question.

We did not assess how well the sequences within the generated blocks align. Methods that explicitly model alignment, such as Minigraph-Cactus, likely produce blocks that are better suited for whole-genome alignment. Nonetheless, future work may combine rearrangement-based and alignment-based synteny block theory. In practice, it may be possible to use the larger blocks generated by MICE as a pre-segmentation step before applying slower alignment-based methods or even seeding aligners with anchors computed by MICE. As of now, both of these possibilities depend on further theoretical work as neither anchors nor partitions generated by MICE are necessarily unique. Additionally, while our results on breakpoints focus on representing rearrangements, we did not evaluate duplication breakpoints due to the challenges of defining this kind of breakpoint (see “Extension to Duplicates” in Section 3.2). This limitation may affect all tools, except for “MICE BP bijection” where we preserve breakpoints by avoiding merging with duplicated elements. Our proposed methods for duplicated elements come with a tradeoff of either more fragmented blocks (“MICE BP bijection”) or losing some theoretical guarantees (“MICE duplicates”). This suggests that a different generalization may yield a more effective handling of duplicates while ensuring stricter guarantees regarding breakpoints. For example, “MICE duplicates” so far guarantees only what is known in the literature as *local collinearity* (collinearity per phrase induced by a block), not *global collinearity* (collinearity w.r.t. the entire genome).

Another research direction is to explore relaxed variants of the theoretical properties presented here. For example, one could require that each pair of genomes shares at least one element in a block, without requiring it to be the same element across all pairs. However, this may come at the risk of a computationally harder problem or too dissimilar blocks. Nonetheless, we observe that even when the optimization problem of finding the smallest synteny block partition may be NP-hard, it could still be practically valuable to find a synteny block partition that retains all biologically motivated constraints (such as breakpoint-freeness) while not guaranteeing to be the smallest. Heuristics in this direction may thus be one way to tackle the NP-hard problems we discussed here. Finally, one could consider the frequency of breakpoints and allow merges across breakpoints that occur only rarely, which may enable stronger compression in closely related genomes, such as pangenome datasets.

## Acknowledgments

This project has received funding from the European Union’s Horizon 2020 research and innovation programme under the Marie Skłodowska-Curie grant agreement number 872539 (PANGAIA). Most of the work was conceived and developed during a research stay of LB and LP at Simon Fraser University. We thank Kamil Hepak for helpful comments on readability, visualization and language.

