## Appendix for "On Deriving Synteny Blocks by Compacting Elements"

#### A Proof for Lemma 1

**Lemma 1.** *Let $S$ be a set of strings over element set $E$, and $P_i$ a contiguous, orientable part of a partition on $E$. If $P_i$ is collinear in $S$, then $P_i$ is breakpoint-free on $S$.*

*Proof.* (By contradiction) Assume for two strings $s, t$, the induced phrases $s^{P_i}$, $t^{P_i}$ have a breakpoint with respect to each other. That is, for some elements $a, b \in p$, $s^{P_i}|_{t^{P_i}}$ contains $a^x b^y$ for some $x, y \in \{h, t\}$ and $t^{P_i}|_{s^{P_i}}$ does not. Without loss of generality, let $x = h$ and $y = t$.

We distinguish two cases: (I) $t^{P_i}|_{s^{P_i}}$ contains $a^h b^t$. (II) $t^{P_i}|_{s^{P_i}}$ contains $a^h c^z$ for some other shared element $c$ with $z \in \{h, t\}$.

In case (I) we have $s^{P_i} = \ldots + aX + b \ldots$ and $t^{P_i} = \ldots + aY - b \ldots$ for substrings of non-shared elements $X, Y$. Then $P_i$ is not orientable.

In case (II) we have $s^{P_i} = A + aX + bB$ and $t^{P_i} = W + aY \pm cC$ for some substrings $A, X, B, W, Y, C$ where elements in $X, Y$ are not shared between $s^{P_i}, t^{P_i}$. Therefore $c$ does not occur in $X$ and $b$ does not occur in $Y$. Since both strings agree on the sign of $a$, we know the partial order should either rise or fall from left to right in both strings (see Def. 2). W.l.o.g. let the partial order rise. We further distinguish three cases.

(i) Let $b$ occur in $W$. Then $t^{P_i}$ implies the order: $b \prec a \prec c$. However, $s^{P_i}$ implies $a \prec b$, a contradiction.

(ii) Let $b$ occur in $C$ and $c$ occur in $A$. Then follows from the order in $s^{P_i}$: $c \prec a \prec b$, which contradicts with $a \prec c$ in $t^{P_i}$.

(iii) Let $b$ occur in $C$ and $c$ occur in $B$. Then in $s^{P_i}$ we have $a \prec b \prec c$ while in $t^{P_i}$ we have $a \prec c \prec b$, a contradiction. □

#### B Proof for Theorem 1

**Theorem 1.** *Let $P$ be an anchored synteny block partition for a set $S$ of singular strings. For each pair of strings $s, t \in S$ there is a bijection $\beta_{s,t}$ between extremities in $\mathrm{Bp}(s,t)$ and in $\mathrm{Bp}(\mathrm{enc}_P(s), \mathrm{enc}_P(t))$, such that*

- *for each breakpoint $\{a, b\} \in \mathrm{Bp}(\mathrm{enc}_P(s), \mathrm{enc}_P(t))$, $\{\beta_{s,t}(a), \beta_{s,t}(b)\}$ is a breakpoint of $s, t$,*
- *for each breakpoint $\{c, d\} \in \mathrm{Bp}(s,t)$, $\{\beta^{-1}_{s,t}(c), \beta^{-1}_{s,t}(d)\}$ is a breakpoint of $\mathrm{enc}_P(s), \mathrm{enc}_P(t)$.*

*Proof.* We first define the mapping $\beta_{s,t}$ for each extremity of a block $P_i$. Let $s^{P_i}, t^{P_i}$ be the phrases induced by $p$ in $s$ and $t$ respectively. Both $s^{P_i}$ and $t^{P_i}$ contain the anchor, therefore they share at least one element. Let $a$ be the first element in $s^{P_i}$ that also occurs in $t^{P_i}$. Because $s^{P_i}$ does not have any breakpoints with $t^{P_i}$, $a$ must either be the first element in $t^{P_i}$ and occur with the same orientation or occur in reverse orientation and as the last shared element of $t^{P_i}$. For the same reason, the last shared element $b$ in $s^{P_i}$ is the last shared element in $t^{P_i}$ or occurs in reverse orientation as the first shared element in $t^{P_i}$. Without loss of generality, $s^{P_i}$ be a forward occurrence of $P_i$. In $\beta_{s,t}$, we then map $i^t$ to the first extremity of $a$ and $i^h$ to the last extremity of $b$.

Consider an adjacency $\{i^x, j^y\}$ in $\mathrm{enc}_P(s)|_{\mathrm{enc}_P(t)}$. Then $\{\beta_{s,t}(i^x), \beta_{s,t}(j^y)\}$ is an adjacency in $s|_t$. To see why, regard $\mathrm{enc}_P(s)$ and $s$. Let $s', s'_{\mathrm{enc}}, t', t'_{\mathrm{enc}}$ be the string obtained by replacing each signed element with its extremities in $s, \mathrm{enc}_P(s), t$ and $\mathrm{enc}_P(t)$ respectively.

We have $s'_{\mathrm{enc}} = \ldots i^x U j^y \ldots$ and accordingly, $s' = \ldots \beta_{s,t}(i^x) V \beta_{s,t}(j^x) \ldots$ for some substrings $U, V$. For the sake of contradiction, assume $V$ contains extremities that occur in $t'$. Then $U$ contains extremities that occur in $t'_{\mathrm{enc}}$. However, then $\{i^x, j^y\}$ is not an adjacency in $\mathrm{enc}_P(s)|_{\mathrm{enc}_P(t)}$. Thus, $\{\beta_{s,t}(i^x), \beta_{s,t}(j^y)\}$ is an adjacency in $s|_t$. An analogous argument holds for adjacencies of $\mathrm{enc}_P(t)|_{\mathrm{enc}_P(s)}$. Since adjacencies are a superset of breakpoints, each breakpoint $\{i^x, j^y\}$ between $\mathrm{enc}_P(s)$ and $\mathrm{enc}_P(t)$ has a corresponding a breakpoint $\{\beta_{s,t}(i^x), \beta_{s,t}(j^y)\}$ between $s, t$.

Now regard a breakpoint $\{a^x, b^y\}$ between $s, t$. W.l.o.g. let $\{a^x, b^y\}$ be an adjacency in $s|_t$, but not in $t|_s$. Then $a, b$ are in different parts of $P$. Otherwise the part would not be breakpoint free or not be contiguous.

Let thus $a, b$ be in different parts $P_i, P_j \in P$. Observe the phrases of $P_i$ and $P_j$ in $s$. Let again $s', s'_{\text{enc}}, t', t'_{\text{enc}}$ be the string obtained by replacing each signed element with its extremities in $s, \text{enc}_P(s), t$ and $\text{enc}_P(t)$ respectively.

Then, $s' = \ldots a^x U_1 U_2 U_3 b^y \ldots$ for some substrings $U_1, U_2, U_3$ of $U$, where the phrase induced by $P_i$ includes $a^x U_1$ and the phrase induced by $P_j$ includes $U_3 b^y$. Since $U = U_1 U_2 U_3$ does not have any extremities that occur in $t'$, $\beta_{s,t}(i^{x'}) = a^x$ and $\beta_{s,t}(j^{y'}) = b^y$ for some $x', y' \in \{h, t\}$. Note that since $U_2$ does not contain any elements in $t$, there are no synteny blocks between $P_i$ and $P_j$ that are also present in $t$. Thus $\{i^{x'}, j^{y'}\}$ is an adjacency in $\text{enc}_P(s)|_{\text{enc}_P(t)}$.

To see why it is a breakpoint, consider that in $t$, $\{a^x, b^y\}$ is not an adjacency in $t|_s$. We distinguish two cases: (I) $\{a^x, b^w\}$ is an adjacency in $t$ for $w \neq y$ and (II) $\{a^x, c^z\}$ is an adjacency in $t|_s$ for some other element $c$ with $z \in \{h, t\}$.

In Case (I), $\text{enc}_P(t)|_{\text{enc}_P(s)}$ contains the adjacency $\{i^{x'}, j^{w'}\}$ with $w' \neq y'$. Therefore, $\{i^{x'}, j^{y'}\}$ is a breakpoint. In Case (II), $c$ must be in a third part $P_l$ as otherwise $P_i$ or $P_j$ would not be breakpoint free. Then $\text{enc}_P(t)|_{\text{enc}_P(s)}$ contains some adjacency $\{i^{x'}, l^{z'}\}$. Therefore, $\{i^{x'}, j^{y'}\}$ is a breakpoint. □

### C Hardness of MLSBP

Here, we show that MLSBP is NP-hard via a reduction from vertex-cover. Let $G = (V, E)$ be a graph and let $r$ be a non-negative integer. The Vertex Cover decision problem asks whether there exists a subset $C \subseteq V$ with $|C| \leq r$ such that every edge in $E$ is incident to at least one vertex in $C$. We describe a polynomial-time reduction from this problem to an instance $S_G$ of our problem MLSBP. For each vertex $v_i \in V$, we construct a gadget represented by a two-character string

$$X_i = xv_i,$$

where $x$ is a common character shared among all vertex gadgets. The signs are always positive, therefore we omit them. For each edge $e_j = (v_p, v_q) \in E$, we construct two gadgets

$$Y_j = \{v_p v_q, v_q v_p\},$$

which introduce a breakpoint between the two corresponding vertex strings. The complete instance is then defined as the union of those gadgets $S_G = X_1 \cup X_2 \cup \ldots \cup X_{|V|} \cup Y_1 \cup Y_2 \cup \ldots \cup Y_{|E|}$. Since each edge gadget has a conflict they need to be separated into two different partitions, therefore into two different phrases each, i.e., $v_p|v_q, v_q|v_p$. Each vertex gadget can be either split into two phrases or kept together. We refer to a vertex where its gadget is split into two phrases as *selected*. With the following theorem we prove that MLSBP is NP-hard.

**Theorem 5.** *Let $G = (V, E)$ be a graph and $r$ a non-negative integer, and let $S_G$ be the set of strings constructed as above. Then $G$ has a vertex cover of size at most $r$ if and only if there exists an MLSBP of $S_G$ such that the total number of phrases is $|V| + 4|E| + r$.*

*Proof.* If two nodes $v_p$ and $v_q$ are connected by an edge, they cannot be placed in the same partition, since they form a breakpoint. Therefore, at least one of the two vertices must be selected. If both are selected, they can be assigned to different partitions, as we aim to minimize the number of phrases and can use as many partitions as needed.

Let $\tau(G)$ denote the size of a minimum vertex cover of $G$. If $\tau(G) \leq r$, we can select $\tau(G)$ vertices in $S_G$, and the total number of phrases is $r' = |V| + 4|E| + \tau(G) \leq |V| + 4|E| + r$. This is because each edge contributes four phrases, and each selected vertex adds one additional phrase. This constitutes a valid solution for $S_G$, since the complement of a vertex cover is an independent set, so no two vertices in it share an edge. Therefore, the character $x$ is in a partition only with characters that do not share an edge.

If $\tau(G) > r$, there are two possible cases: (1) if we select more than $r$ vertices, it is clear that $r' > |V| + 4|E| + r$; (2) if we select at most $r$ vertices, there remain $\tau(G) - r$ vertices that still share an edge, and therefore cannot be placed in the same partition as the character $x$, leading to a contradiction. □

#### D Hardness of MSSBP

In addition to MLSBP, MSSBP is also NP-hard. We show the reduction from the Boolean satisfiability problem (SAT).

In short, SAT asks whether a formula $F$ in propositional logic can be satisfied. This formula is built over an alphabet of variables $V$, utilizing each $x \in V$ as *literals*, i.e., as "positive" $x$ or "negative" $\neg x$. These literals are connected with logical operators $\vee$ ("or") and $\wedge$ ("and"). Each variable can be assigned a Boolean value, i.e., "true" or "false".

The formula $F$ is given in *conjunctive normal form*, that is $F = c_1 \wedge c_2 \ldots \wedge c_m$ where each *disjunctive clause* $c_i$ consists of disjunctions of variable literals $c_i = l_i^1 \vee l_i^2 \ldots \vee l_i^{m_i}$, where each $l_i^j = x$ or $l_i^j = \neg x$ for some variable $x \in V$. The question asked in SAT is whether there is an assignment of Boolean values ("true", "false") to each variable $x$, such that $F$ is true. SAT is NP-complete.

**Theorem 6.** *For any given SAT formula $F$ in CNF with $n$ variables and $m$ clauses, there is an instance of MSSBP with size $\mathcal{O}(nm + n^2)$, such that the smallest synteny block partition has size $2n$ if and only if $F$ is satisfiable. All synteny blocks in this partition are then also collinear.*

*Proof.* In this proof, all elements have positive orientation. For the sake of readability, we thus omit their (positive) signs.

Note that we can ensure that two elements $e, d$ are in different partitions by adding the chromosomes $ed$ and $de$. We write $\mathtt{DIFF}(d, e) = \{de, ed\}$.

For each variable $x_i$, we add three elements, namely $\mathtt{t}_i, \mathtt{f}_i, \mathtt{v}_i$. We model the boolean value of each variable $x_i$ by grouping $\mathtt{v}_i$ either with $\mathtt{t}_i$ ("true") or $\mathtt{f}_i$ ("false"). To ensure a variable is either true or false, we add the chromosomes $\mathtt{DIFF}(\mathtt{t}_i, \mathtt{f}_i)$. For each pair of different variables $x_i \neq x_j$, we add the chromosomes $\mathtt{DIFF}(a, b)$ for all $a \in \{\mathtt{t}_i, \mathtt{f}_i, \mathtt{v}_i\}$ and all $b \in \{\mathtt{t}_j, \mathtt{f}_j, \mathtt{v}_j\}$ to ensure distinct parts for each variable. Note that there are now at least $2n$ parts in any synteny block partition.

For each disjunctive clause $c_i$, we add an element $\mathtt{c}_i$. For each clause $c_i$ and variable $x_j$, we add $\mathtt{t}_j\mathtt{v}_j\mathtt{c}_i$ if variable $x_j$ occurs as a positive literal $x_j$ in $c_i$ and $\mathtt{DIFF}(\mathtt{t}_j, \mathtt{c}_i)$ otherwise. We add $\mathtt{c}_i\mathtt{v}_j\mathtt{f}_j$ if variable $x_j$ occurs as a negative literal $\neg x_j$ in $c_i$ and $\mathtt{DIFF}(\mathtt{f}_j, \mathtt{c}_i)$ otherwise. In cases where a disjunctive clause contains both $x_j$ and $\neg x_j$, we omit the clause as it is automatically satisfied by any Boolean assignment.

Note that if a clause element $\mathtt{c}_i$ is grouped with $\mathtt{t}_j$ (or $\mathtt{f}_j$), due to contiguousness $\mathtt{v}_j$ is also in $\mathtt{t}_j$ (or $\mathtt{f}_j$), i.e., the clause is satisfied by the corresponding boolean assignment to $x_j$. If there is a partition with $2n$ parts, we can thus derive a satisfying assignment of boolean values to all $x_j$.

Conversely, given a satisfying assignment to $F$, we can obtain a partition by grouping all $\mathtt{v}_j$ with their corresponding boolean value and each $\mathtt{c}_i$ with any variable that satisfies it, obtaining a partition of the minimum size $2n$.

Note that in each case, we have the partial order $\mathtt{t}_i \prec \mathtt{v}_i \prec \mathtt{c}_j$ for $\mathtt{c}_j$ being grouped with $\mathtt{t}_i$ or $\mathtt{c}_j \prec \mathtt{v}_i \prec \mathtt{f}_i$ for $\mathtt{c}_j$ being grouped with $\mathtt{f}_i$ that satisfies Definition 2. The blocks are thus collinear.

Moreover, the MSSBP instance is comprised of $\mathcal{O}(nm + n^2)$ strings, each with at most 3 elements. The size of the MSSBP instance is thus $\mathcal{O}(nm + n^2)$. □

#### E Constructing Anchored Collinear Blocks by Unions

**Lemma 5.** *Given an anchored, collinear synteny block $P_j = \{e_1, \ldots e_m\}$ with anchor $c$ and its partial order $\prec$. Let $P_\prec = \{e \in P_j \mid e \prec c\}$ and $P_\succ = \{e \in P_j \mid e \succ c\}$. Let $Q_0 = P_\prec \cup \{c\}$ and $Q_{i+1} = Q_i \cup \{q_{i+1}\}$ for a topological sorting $q_1 \ldots q_l$ of $P_\succ$. Then each $(Q_i)_{i=0}^{l}$ is anchored, collinear and $\{q_{i+1}\} \to Q_i$ for all $0 \leq i < l$.*

*Proof.* $Q_i$ is collinear and has anchor $c$ because $P_j$ is collinear and has anchor $c$.

Regard $q_{i+1}$. Let $s$ be a string in which $q_{i+1}$ occurs. Since $c$ is an anchor, $s$ also contains $c$. W.l.o.g. let $c$ occur before $q_{i+1}$ (otherwise simply invert the string). Then $s = \ldots cT \pm q_{i+1} \ldots$ for some substring $T$. Then all elements in $T$ must be in $P$, otherwise $P$ would not be contiguous in $s$. Additionally, for each element $t$ in $T$ must hold $t \prec q_{i+1}$. Thus for each element $t$ in $T$, $t \in Q_i$. The same argument holds for all strings where $q_{i+1}$ is present. Additionally, since $P_j$ is orientable, $q_{i+1}$ always occurs in the same orientation relative to $c$. Thus, if $Q_i$ is contiguous, $\{q_{i+1}\} \to Q_i$.

We now show contiguousness by induction. $Q_0$ is contiguous, otherwise $P_j$ would not be contiguous or not collinear. Assume $Q_i$ is contiguous. Then $\{q_{i+1}\} \to Q_i$. Thus, using Lemma 2, $Q_{i+1} = Q_i \cup \{q_{i+1}\}$ is contiguous. □

**Proposition 1.** *Any collinear, anchored synteny block* $P_j = \{e_1, \dots e_m\}$ *on string set* $S$ *and element set* $E$ *can be constructed by iteratively unifying pairs of parts* $R_i, R_l$ *where* $R_i \to R_l$ *from some partition* $\{\{e_1\}, \dots, \{e_m\}\} \cup R$ *of* $E$.

*Proof.* Let $\prec$ be the partial order on $P_j$. Consider the two sets $P_\prec = \{x \in P \mid x \prec c\}$ and $P_\succ = \{x \in P \mid x \succ c\}$. Note that since $c$ is an anchor, $P_\prec, P_\succ$ are disjoint.

Regard the set $Q = P_\prec \cup \{c\}$. It is contiguous and collinear because $P$ is.

Using an inverted partial order $\prec'$, i.e., $x \prec' y \iff y \prec x$, we can then construct $Q$ with unions as described in Lemma 5 starting with $Q_0 \coloneqq \{c\}$.

We can then construct $P$ with unions as described in Lemma 5 starting with $Q_0 \coloneqq Q$. □

Since we can construct each part in this manner, this proves Theorem 3.

#### F Linear Time and Space Algorithm

To keep track of the partitions formed, we maintain an array MERGEDTO of size equal to the number of distinct elements, where MERGEDTO$[e]$ stores the element into which element $e$ was merged, initialized as MERGEDTO$[e] = e$ for all $e \in E$.

We store three main data structures. Each input string is represented as a linked list of occurrences of extremities. Let $e_i^x$ denote the $i$-th occurrence of an extremity $e^x$ in any of the input strings. For every occurrence $e_i^x$, we store the extremity adjacent to it as

$$\text{NEXT}(e_i^x) = f_j^y.$$

If $e_i^x$ occurs at the end of a string, we set $\text{NEXT}(e_i^x) = \$$, where \$ denotes the telomere.

In addition, for each extremity $e^x$ we maintain the list of its occurrences,

$$\text{OCC}(e^x) = [e_1^x, e_2^x, \dots],$$

which allows us to access all instances of that extremity across the input. The two extremities of the same element always have occurrence lists of the same length, and the $i$-th entry of $\text{OCC}(e^t)$ corresponds to the same copy of the element as the $i$-th entry of $\text{OCC}(e^h)$.

In addition, we also store for each extremity $e^x$ the collection of extremities that appear adjacent to it in any of the input strings. We denote it by $\text{ADJ}(e^x)$, and store it as a vector. Initially, it contains all extremities adjacent to $e^x$ in any of the strings.

$$\text{ADJ}(e^x) = [f^y, g^z, \dots,].$$

An extremity $e^x$ has a unique neighbor if all of the extremities in $\text{ADJ}(e^x)$ are the same.

We now prove that Algorithm 1 can be implemented in linear time and space. The complexity analysis consists of three parts: how many times the main while loop is executed, the cost of reconnecting adjacent values after merging one element with another, and the cost of checking whether any extremity after the update has a unique neighbor.

The while loop of Algorithm 1 is implemented as a queue that is processed until it becomes empty. Initially, the queue contains all extremities that have a unique neighbor. During each iteration, one extremity is removed from the queue, check if it still has a unique neighbor, and if the corresponding merge creates a new unique neighbor, it is added to the queue. Each adjacency between two extremities is considered for merging only once, and at most two extremities can be inserted into the queue for the same adjacency, in the case where each is the unique neighbor of the other. Therefore, the total number of queue insertions, and consequently the number of iterations of the while loop, is bounded by twice the number of adjacencies, which is linear in the total number of extremity occurrences $L$.

When the queue pops an extremity $a^x$ with its unique adjacent extremity $b^y$, the element $a$ is merged into $b$. Let $a^{\bar{x}}$ denote the opposite extremity of $a^x$,i.e., $\bar{x} = t$ if $x = h$, and $\bar{x} = h$ if $x = t$. All extremities that were adjacent to $a^{\bar{x}}$ must now be connected to $b^y$. To perform this update, we inspect every occurrence $a_i^{\bar{x}} \in \textsc{occ}(a^{\bar{x}})$, and for each $\textsc{next}(a_i^{\bar{x}}) = c_l^z$, we append $c^z$ to $\textsc{adj}(b^y)$ and $b^y$ to $\textsc{adj}(c^z)$. This is done for every single occurrence of $a_i^{\bar{x}}$, even if it implies adding multiple times $c^z$ to $\textsc{adj}(b^y)$. At the same time, we update the linked list to preserve the structure of the strings: for each occurrence $a_i^x \in \textsc{occ}(a^x)$ we update the corresponding $b_j^y$ adjacent to $a_i^x$, and we set $\textsc{next}(b_j^y) = c_l^z$. The same we do for each occurrence of $c_l^z$, by setting $\textsc{next}(c_l^z) = b_j^y$. This operation is performed exactly once for each occurrence of element $a$. Since removed elements are never reinserted, the total amortized cost of these updates is $\mathcal{O}(L)$. Moreover, the adjacency lists $\textsc{adj}(c^z)$ and of $\textsc{adj}(b^y)$ are updated simply by appending $b^y$ and $c^z$, respectively, to the end of the lists.

Lastly, we need to check whether any extremities has now a unique neighbor. This must be done for each $c^z$ and for $b^y$. To perform this efficiently, we use a function IsUniqueNeighbor defined in Algorithm 2, which takes in input a list of adjacencies $\textsc{adj}(e^x)$ and returns true if the there is only one adjacent extremity or false otherwise. This function scans $\textsc{adj}(e^x)$ until it finds an extremity $e^x$ which was not merged yet. This can be checked in constant time by verifying that $\textsc{mergedto}(e) = e$. If such an extremity $f^y$ is found, the function continues scanning for another non-merged extremity different from $f^y$. If a second one is found, or if all elements have already been merged, the function returns false; otherwise, it returns true. We also maintain an auxiliary data structure that stores the index of the first non-merged extremity, which we refer to as $\textsc{adjptr}(a^x)$. When two distinct non-merged neighbors are discovered, they are moved next to each other in the adjacency list so that subsequent checks can be performed in constant time.

**Algorithm 2** Check if an extremity $e^x$ has a unique neighbor.

**procedure** IsUniqueNeighbor($e^x$, ADJ, ADJPTR, MERGEDTO)
    $i \leftarrow \textsc{adjptr}(e^x)$
    $f^y = \textsc{adj}(e^x)[i]$
    **while** $i \leq |\textsc{adj}(e^x)| \wedge \textsc{mergedto}[e] \neq e$ :
        $i \leftarrow i + 1$
        $f^y = \textsc{adj}(e^x)[i]$
    **if** $i \leq |\textsc{adj}(e^x)|$ :
        $g^z = \textsc{adj}(e^x)[i]$
        **while** $i \leq |\textsc{adj}(e^x)| \wedge (\textsc{mergedto}[g] \neq g) \vee g^z = f^y)$ :
            $i \leftarrow i + 1$
            $g^z = \textsc{adj}(e^x)[i]$
        $\textsc{adjptr}(e^x) \leftarrow i$
        $\textsc{adj}(e^x)[i] \leftarrow f^y$
        **return** $\textsc{adjptr}(e^x) = |\textsc{adj}(e^x)|$
    **return false**

This function is called once for each occurrence of an element $a$ which is merged, plus once for the element $b$ into which $a$ is merged. Each call takes either constant time or increases the scanning index monotonically until one or two non-merged elements are found. Since the total size of all adjacency lists ADJ is initially $L$, and each merge operation appends to these lists once per occurrence of an element, the total size remains $\mathcal{O}(L)$. Therefore, IsUniqueNeighbor runs in amortized $\mathcal{O}(L)$ time.

For similar reasons, the space used is linear in $L$.

#### G Integrating Duplicates

##### G.1 Generalizing the problem definition

We have seen that a synteny block can be defined for singular strings as a set of elements that induces at most one contiguous phrase per string. When involving duplicates, this definition is no longer viable. Instead, we will view a synteny block $R_i$ on a set of strings $S = \{s_1, \ldots, s_n\}$ as a set of phrases or more accurately phrase indices $R_i \subset \{(j, l, r) \mid 1 \leq j \leq n, 1 \leq l \leq r \leq |s_j|\}$, i.e., if $(j, l, r) \in R$, then $s_j[l : r]$ is one of the phrases of block $R$.

**Definition 6.** *A set of phrases* $R_i \subset \{(j,l,r) \mid 1 \leq j \leq n, 1 \leq l \leq r \leq |s_j|\}$ *with* induced part $P_{R_i} := \bigcup_{(j,l,r)\in R} \mathrm{elm}(s_j[l : r])$, *is a* synteny block *on strings* $S = \{s_1, \ldots, s_n\}$ *with element set* $E$, *if all of the following hold:*

1. *Phrases* $(j,l,r) \neq (j,l',r')$ *in the same string* $s_j$, *do not overlap, i.e.,* $l \leq r < l' \leq r'$ *or* $l' \leq r' < l \leq r$.
2. *For each element* $e \in P$ *and occurrence of* $e$, $s_j[x] = e$, *there is a phrase of* $R_i$ *that contains it, i.e.,* $\exists (j,l,r) \in R$ *with* $l \leq x \leq r$.
3. $R_i$ *is* orientable, *i.e., there is an orienting function* $f$, *such that each element occurs only with one orientation in* $\{f(s_j[l : r])s_j[x] \mid (j,l,r) \in R, l \leq x \leq r\}$.
4. *For each* $(j,l,r) \in R_i$, *the associated phrase* $s_j[l : r]$ *is a singular string.*
5. *The number of occurrences of* $R_i$ *per string* $s_j$ *is equal to the number of occurrences of some element* $e \in P_{R_i}$ *in* $s_j$ $|\{(l,r) \mid (j,l,r) \in R\}| = |\{x \mid \mathrm{elm}(s_j[x]) = e\}|$.
6. $R_i$ *is* breakpoint free, *i.e., any two phrases* $s_j[l : r], s_{j'}[l' : r']$ *for* $(j,l,r),(j',l',r') \in R_i$ *have no breakpoints with respect to each other.*

In this definition, Items 1 and 2 ensure $R$ behaves like a part in a partition, Item 3 is equivalent to orientability, Items 4 and 5 ensure a version of contiguousness for duplicates and Item 6 ensures breakpoint freeness. Observe that for singular strings, Definition 6 is equivalent to synteny blocks described in Definition 1. We can then define a synteny block partition as follows.

**Definition 7.** *A collection of synteny blocks* $R = \{R_1, \ldots, R_m\}$ *with induced parts* $P = \{P_{R_1}, \ldots, P_{R_m}\}$ *on string set* $S$ *on elements* $E$ *is a* synteny block partition *if* $P$ *is a partition of* $E$.

The parsing of $S$ induced by $R$ then also defines an encoding of the strings in $S$ as strings on the alphabet $\{1, 2, \ldots, m\}$ where, for every string $s \in S$, its encoding $\mathrm{enc}_R(s)$ is obtained from $s$ by replacing every phrase of $s$ in $R_i$ with $+i$ (resp. $-i$) if $R_i$ is a forward (resp. backward) occurrence in $s$.

We can then analogously to Problems 1 and 2 define two new problems.

*Problem 5.* Given a collection of strings $S = \{s_1, \ldots, s_m\}$ on the element set $E$ and a set $\mathcal{C}$ of constraints on signed partitions of $E$, find a synteny block partition $R$ for $S$ such that $R$ satisfies all constraints in $\mathcal{C}$, and

[*Minimum-Length Synteny Blocks Problem* (MLSBP)] $\sum_{i=1}^{m} |\mathrm{enc}_R(s_i)|$ is minimized;
[*Minimum-Size Synteny Blocks Problem* (MSSBP)] the size of the partition $|R|$ is minimized.

Note that singular strings are a special case of arbitrary strings. The general formulations of MLSBP and MSSBP are thus NP-hard for arbitrary strings as well. The definitions for collinearity and anchored blocks generalize as well.

**Definition 8 (Collinear).** *Let* $S$ *be a set of strings. A synteny block* $R_i$ *with induced part* $P_i$ *and orienting function* $f$ *is* collinear *in* $S$ *if there exists a partial order* $\prec$ *on the signed elements in* $P_i$ *such that, for every* $(k,l,r) \in R_i$:

$$\forall\ l \leq j < r,\ \textit{if } f(s_k[l : r]) = +,\ \textit{then } s_k[j] \prec s_k[j+1],\quad \textit{otherwise } [f(s_k[l : r]) = -] s_k[j+1] \prec s_k[j].$$

Lemma 1 also holds for this definition of collinearity. The proof is analogous to that in Lemma 1.

**Definition 9 (Anchored).** *Let* $S$ *be a set of strings and* $R = \{R_1, \ldots, R_k\}$ *a synteny block partition for* $S$. *We say that* $R_i$ *is* anchored *if* $\bigcap_{(j,l,r)\in R_i} \mathrm{elm}(s_j[l : r])$ *is not empty. We call any* $e \in \bigcap_{(j,l,r)\in R_i} \mathrm{elm}(s_j[l : r])$ *an* anchor *of* $R_i$. *We say that* $R$ *is anchored if all synteny blocks* $R_i$ *are anchored.*

We show in the next subsection how to generalize the algorithm for synteny block construction.

##### G.2 Generalizing the Algorithm

The concept of unique neighbors is the same for duplicates. Just note that now in a single string, an extremity can have multiple neighbors. For example regard $+a$ $+b$ $+a$ $+c$. In this string, we have $b^t \to a^h$, but not $a^h \to b^t$, since there is another adjacency $a^h c^t$.

**Lemma 6.** *Let $R_a$ and $R_b$ be two anchored, collinear synteny blocks of the same synteny block partition and let $R_a \to R_b$. Let $c_a, c_b$ be the anchors of $R_a, R_b$ respectively.*

*Then there is a set of intervals $R_a \sqcup R_b$ that covers the same positions as $R_a$ and $R_b$ and that is a collinear synteny block with anchor $c_b$.*

*Proof.* W.l.o.g. let $a^h \to b^t$. Then for each forward block $(j, l_a, r_a) \in R_a$, there is a forward block $(j, r_a + 1, r_b) \in R_b$. Analogously, for each backward block $(j', l'_a, r'_a) \in R_a$, there is a backward block $(j', l'_b, l'_a - 1) \in R_b$. We can then extend these blocks as $(j, l_a, r_b)$ or $(j', l'_b, r'_a)$ respectively. $R_a \sqcup R_b$ is thus defined as all phrases of $R_b$, each extended in this manner with at most one phrase from $R_a$.

The positions covered are then the same as for $R_a$ and $R_b$ separately and each phrase contains element $c_b$. The part induced by $R_a \sqcup R_b$ is then $P_a \cup P_b$ where $P_a, P_b$ are the induced parts of $R_a, R_b$ respectively. A new partial order is obtained by concatenating the partial orders of $R_a, R_b$.

The proofs for other $x, y \in \{h, t\}$ are follow by choosing different orienting functions for $R_a, R_b$. □

Again, we can prove the following theorem, which is analogous to Theorem 3 in the main text. To that end, let $\text{Pos}(e) = \{(j, i, i) \mid \text{elm}(s_j[i]) = e\}$.

**Theorem 7.** *Any collinear anchored synteny block partition $R = \{R_1, \ldots, R_m\}$ on element set $E$ can be constructed by iteratively unifying ($\sqcup$) pairs of blocks $X, Y$ where $X \to Y$ from the base set of blocks $\{Pos(e) \mid e \in E\}$.*

The proof follows the same strategy as in Section 3.2, starting with an analogue to Lemma 5.

**Lemma 7.** *Given a synteny block $R$ with $R = \{(k_1, l_1, r_1), \ldots, (k_m, l_m, r_m)\}$ with anchor $c$ and its partial order $\prec$. Let $P$ be its induced part and $P_\prec = \{e \in P \mid e \prec c\}$ and $P_\succ = \{e \in P \mid e \succ c\}$. Let $q_1, \ldots q_l$ be a topological sorting of $P_\succ$. Let $x_{i,j}$ be the position of character $q_i$ in phrase $(k_j, l_j, r_j)$ if it exists. Let $c_j$ be the position of $c$ in phrase $j$. Let $(k_j, l^0_j, r^0_j) = (k_j, l_j, c_j)$ if the partial order rises in the prase and $(k_j, l^0_j, r^0_j) = (k_j, c_j, r_j)$ if the partial order falls in the phrase. Let $Q_i = \{(k_j, x_{i,j}, x_{i,j}) \mid$ if $q_i$ occurs in phrase $j\}$ and $R_0 = \{(k_j, l^0_j, r^0_j) | 1 \le j \le m\}$.*

*Then $Q_i \to R_i \quad \forall 1 \le i \le l$ if $R_{i+1} = R_i \sqcup Q_i$. Additionally, each $R_i$ is collinear.*

*Proof.* Observe that $R_0$ is a collinear synteny block with anchor $c$ and part $P_\prec \cup \{c\}$. Let $q_i$ occur in phrase $j$. Assume the partial order rises for phrase $j$. Let the phrase in $R_i$ be $(k_j, l^i_j, r^i_j)$. Since the partial order rises $x_i > r^i_j$. For the sake of contradiction, assume $r^i_j + 1 < x_i$. Let the element at that position be $p = s_k[r^i_j + 1]$. Then $p \prec q_i$. However then $p$ must already be in block $R_i$, a contradiction. Therefore $q_i$ always has a neighbor from $R_i$ when the partial order is rising. An analogous argument can be made for falling partial order. Thus, each phrase of $Q_i$ is always preceded or succeeded by elements of $R_i$, $Q_i \to R_i$. Thus, according to Lemma 6, $R_{i+1} = R_i \sqcup Q_i$ is a collinear block with anchor $c$.

□

**Proposition 2.** *Any collinear, anchored synteny block $R$ on element set $E$ can be constructed by iteratively unifying pairs of blocks $X, Y$ where $X \to Y$ from the base set of blocks $\{Pos(e) \mid e \in E\}$.*

*Proof.* The proof is analogous to that of Proposition 1 and uses Lemma 7. □

Again, since any synteny block can be constructed in this manner, the synteny block partition can be constructed. This concludes the proof Theorem 7. With a slight modification, we can then obtain an algorithm for strings with duplicates. We give this modified algorithm as Algorithm 3.

Again, we have similar observations to Observations 1 and 2 and Lemma 3.

**Observation 3** *Let $R = \{R_1, \ldots, R_k\}$ be a collinear anchored synteny block partition of element set $E$ with canonical anchors $c_1, \ldots c_k$ on string set $S$. Then the string set $S'$ where each element but $c_1, \ldots c_k$ is removed and then each $+c_i$ $(-c_i)$ is replaced by $+i$ $(-i)$ is an encoding for $R$.*

**Algorithm 3** Generate a partition $R$ of collinear, anchored synteny blocks and its encoding $S^*$.

**Require:** A set of signed strings $S$ on element set $E = \{e_1, \dots e_l\}$
  $R \leftarrow \{\{\text{Pos}(e_1)\}, \dots, \{\text{Pos}(e_l)\}\}$
  **while** $R_i \to R_j$ for some $R_i, R_j \in R$ with canonical anchors $c_i, c_j$ :
    $R \leftarrow R \setminus \{R_i, R_j\} \cup \{R_i \sqcup R_j\}$
    $r \leftarrow c_i$
    **if** $R_j \to R_i$ : ▷ If they are each other's unique neighbor, remove the larger core
      $r \leftarrow \max(c_i, c_j)$
    Remove $r$ from $S$
  Obtain $S^*$ by replacing each $c_i$ with $i$.
  **return** $R, S^*$

**Observation 4** *Given a set of strings $S = \{s_1, \dots, s_m\}$ and a collinear anchored partition $R = \{R_1, \dots, R_k\}$, let $S' = \{s'_1, \dots s'_m\}$ be the strings obtained by removing an element $e$ that is not a canonical anchor. Let $R' = \{R'_1, \dots R'_k\}$ be the set of sets of phrases, where the coordinates of each phrase $R_i$ have mapped to the coordinates of $S'$ Then $R'$ is a collinear anchored synteny block partition on $S'$.*

**Lemma 8.** *For two elements $e, d$ with $e \to d$. If $d \not\to e$ or $d < e$ and $e$ is a canonical anchor in an anchored, collinear synteny block partition $R = \{R_1, \dots, R_k\}$, then there is a shorter synteny block partition $R'$ with a shorter encoding than $R$, such that $R'$ is also anchored and collinear.*

*Proof.* (By contradiction) Assume that $e$ is a canonical anchor of block $R_i$. Then $d \notin P_{R_i}$. Let $d$ be in block $R_j$ with canonical anchor $c$. Then $R_i \to R_j$ (because $e \to d$), that is $R_i \sqcup R_j$ is a collinear synteny block and has anchor $c$ according to Lemma 6. Observe that $R' = R \setminus \{R_i, R_j\} \cup \{R_i \sqcup R_j\}$ has both fewer parts and a shorter encoding than $R$. □

We arrive at the following corollary.

**Corollary 2.** *Let $S^*$ be the string set produced by Algorithm 3.*

*Let $R^*_{Size}$, $R_{Size}$ be the collinear, anchored partitions on $S^*$ and on $S$ respectively, such that $|R^*_{Size}|$ and $|R_{Size}|$ are minimized. Then $|R^*_{Size}| \leq |R_{Size}|$.*

*Let $R^*_{Length}, R_{Length}$ be collinear, anchored partitions on $S^*$ and on $S$ respectively, such that the encoding lengths $p^* := \sum_{s^* \in S^*} |\text{enc}_{R^*_{Length}}(s^*)|$ and $p := \sum_{s \in S} |\text{enc}_{R_{Length}}(s)|$ are minimized. Then $p^* \leq p$.*

Again, there are no further unique neighbors on $S^*$, therefore $R$ is as large as the optimal solution on $S^*$ and thus $R$ is optimal.

**Theorem 8.** *Algorithm 3 constructs a signed partition $P$ that is a solution to the collinear, anchored MLSBP and collinear anchored MSSBP for general strings.*

We note however, that while each individual phrase constructed by the algorithm is breakpoint free to any other phrase (due to collinearity), breakpoints between unique elements can still be obscured by this procedure. We give an example in Figure 7.

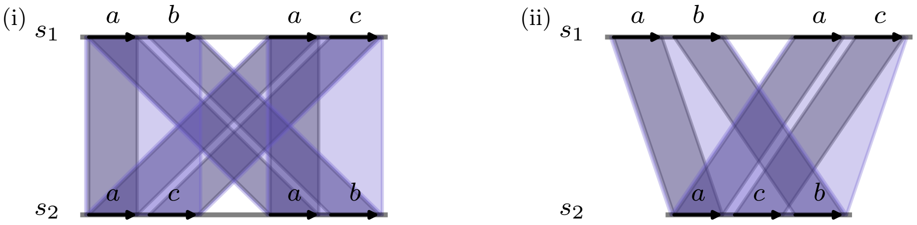


**Fig. 7.** Breakpoints obscured by duplicates. In each case there is a breakpoint between $b$ and $c$ when projecting to singular and shared elements. However, even though the synteny block partition $R$ with a single induced part $\{a, b, c\}$ shown here is breakpoint-free, $R$ obscures this breakpoint. In (i), the encoding of both strings is the same, $\text{enc}_R(s_1) = \text{enc}_R(s_2) = +1 + 1$. The difference in element order between both strings cannot be retrieved from the encoding alone. Here, the rearrangement is still recoverable by disambiguating the encoding using the original element information, i.e. $s_1 \approx +1_1 + 1_2, s_2 \approx +1_2 + 1_1$. In (ii), $\text{enc}_R(s_1) = +1 + 1$ and $\text{enc}_R(s_2) = +1$; the breakpoint is also lost. In this case, one cannot retrieve the breakpoint by disambiguating the encoding of the blocks.

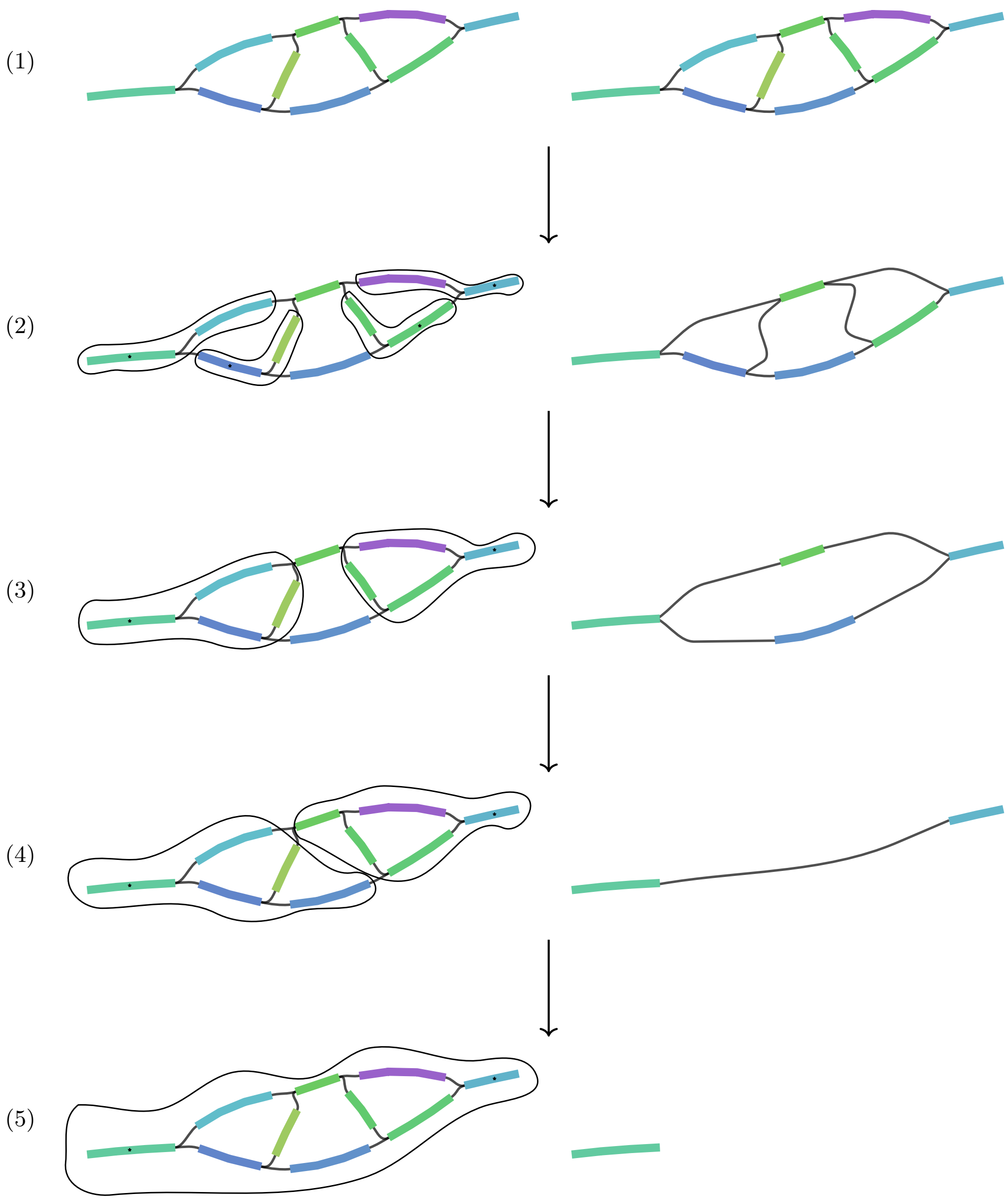


**Fig. 8.** Illustration of Algorithm 1 and 3 on a complex bubble in a pangenome graph. Left: Each part with more than one element is circled. Anchor elements are marked by an asterisk. Right: The graph for modified $S$ where only canonical anchor elements remain is shown. In Illustrations (1) to (5), each new part is constructed as the union $P_i \cup P_j$ of two parts $P_i, P_j$ in the previous illustration, such that $P_i \rightarrow P_j$. Note that such a union may only be one choice among several possible choices. Also note that going from one illustration to the next requires several iterations of the while loop.

#### H Additional Evaluation

**Table 1.** Benchmarking datasets used in Section 4. Of the 16 *Mus musculus* assemblies, only the first four were used for experiments other than the runtime benchmark.

| Taxon | Accession.Version |
|---|---|
| *Y. pestis* | GCF_000006645.1, GCF_000007885.1, GCF_000009065.1, GCF_000013805.1, GCF_000013825.1, GCF_000016445.1, GCF_000018805.1, GCF_000022805.1, GCF_000022845.1, GCF_000186725.1, GCF_000222975.1, GCF_000590535.2, GCF_000834235.1, GCF_000834275.1, GCF_000834315.1, GCF_000834335.1, GCF_000834495.1, GCF_000834755.1, GCF_000834775.1, GCF_000834825.1, GCF_000834845.1, GCF_000834885.1, GCF_000834925.1, GCF_000834985.1, GCF_001188695.1, GCF_001188715.1, GCF_001188755.1, GCF_001188775.1, GCF_001188795.1, GCF_001188815.1, GCF_001188935.1, GCF_001293415.1, GCF_001693595.1, GCF_003798205.1, GCF_003798225.1, GCF_003798345.1, GCF_009295925.1, GCF_009295945.1, GCF_009295965.1, GCF_009295985.1, GCF_009296005.1, GCF_009363195.1, GCF_015159615.2, GCF_015190655.1, GCF_015336085.1, GCF_015336265.1, GCF_015336465.1, GCF_015336695.1, GCF_015336865.1, GCF_015337085.2, GCF_015337285.1, GCF_015337445.1, GCF_015337645.1, GCF_015337825.2, GCF_015338045.1, GCF_015338205.1 |
| *E. coli* | GCF_001612475.1, GCF_001645235.2, GCF_001890265.1, GCF_002057245.1, GCF_002249955.1, GCF_003017915.1, GCF_003017935.1, GCF_003018035.1, GCF_003018055.1, GCF_003018555.1, GCF_003018815.1, GCF_003018835.2, GCF_003018995.1, GCF_003203755.1, GCF_003308955.1, GCF_003308975.1, GCF_003571685.1, GCF_003966445.1, GCF_003966465.1, GCF_004010615.1, GCF_004010655.1, GCF_004358365.1, GCF_004358405.1, GCF_005221585.1, GCF_005221605.1, GCF_005221645.1, GCF_005221685.1, GCF_005221725.1, GCF_005221745.1, GCF_005221785.1, GCF_005221825.1, GCF_005221865.1, GCF_005221985.1, GCF_005222065.1, GCF_005222265.1, GCF_005886035.1, GCF_006874785.1, GCF_006965465.1, GCF_008761535.2, GCF_008925965.1, GCF_008931135.1, GCF_009432415.1, GCF_011330935.2, GCF_014295215.1, GCF_014295315.1, GCF_016776325.1, GCF_019915525.1, GCF_900635325.1 |
| *S. cerevisiae* | GCA_000146045.2, GCA_001051215.1, GCA_001580425.1, GCA_002885995.1, GCA_003086655.1, GCA_003709285.1, GCA_004014915.1, GCA_004328465.1, GCA_018219195.1, GCA_021172205.1, GCA_022626425.2, GCA_022695735.1, GCA_023508825.1, GCA_024732265.1, GCA_024972935.1, GCA_024972955.1, GCA_030292175.1, GCA_030607045.1, GCA_041294695.1, GCA_045517165.1, GCA_051861825.1, GCA_903819125.2, GCA_903819135.2, GCA_903819145.2, GCA_903819155.2, GCA_903819175.2, GCA_903819185.2, GCA_903819195.2, GCA_903819205.2 |
| *A. thaliana* | GCA_023115395.1, GCA_028009825.2, GCA_051624255.1, GCA_051624265.1, GCA_946409825.1 |
| *M. musculus* | GCA_000001635.8, GCA_001624185.1, GCA_001624215.1, GCA_001624295.1, GCA_001624445.1, GCA_001624475.1, GCA_001624505.1, GCA_001624535.1, GCA_001624675.1, GCA_001624745.1, GCA_001624775.1, GCA_001624835.1, GCA_001632525.1, GCA_001632555.1, GCA_001632575.1, GCA_001632615.1 |

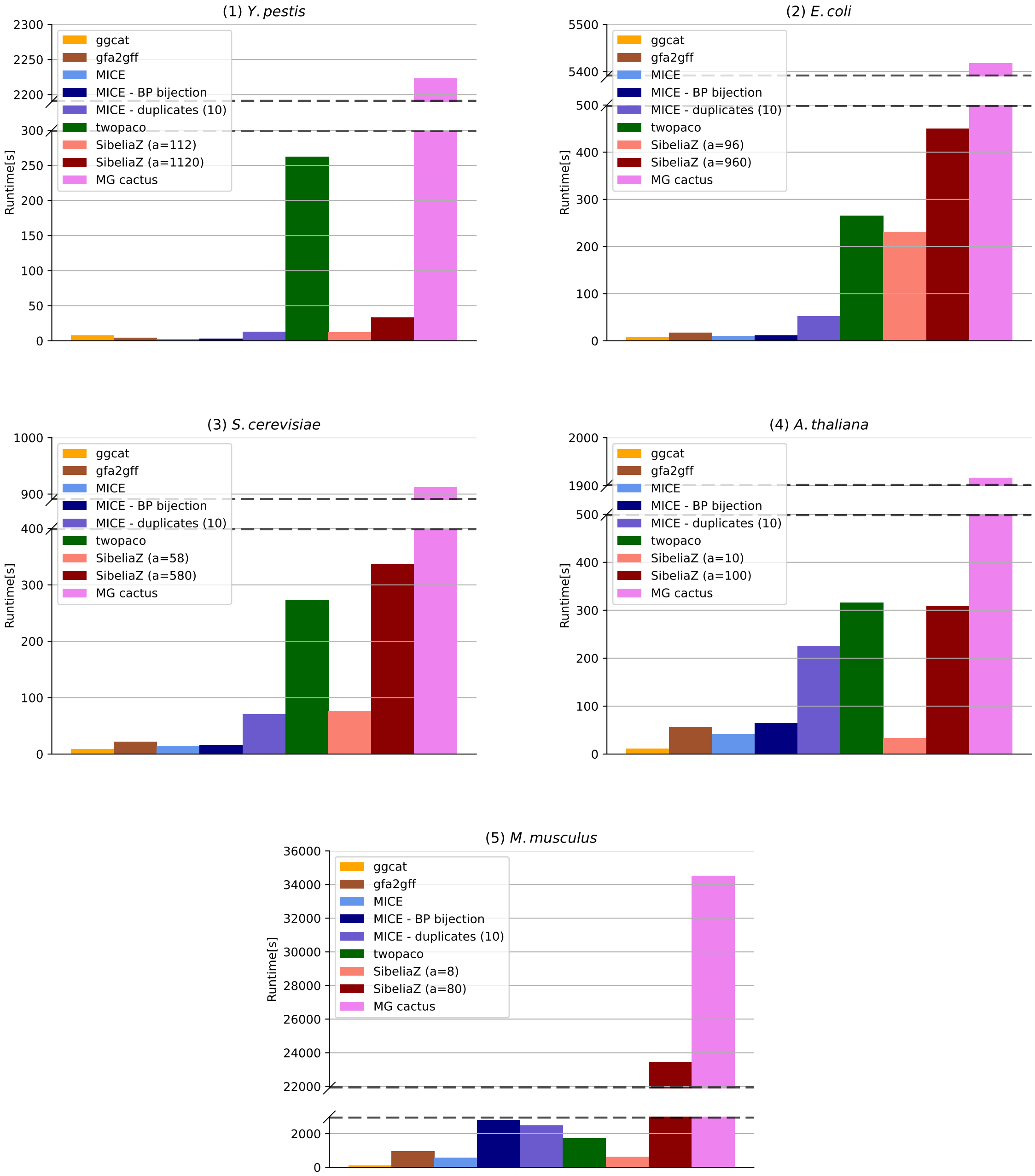


**Fig. 9.** Runtimes of all synteny block generation steps. In this evaluation we used only 4 of the 16 assemblies of the *M. musculus* dataset. Note that the runtimes are generally not comparable, as we ran preprocessing steps, i.e., twopaco for SibeliaZ and ggcat and gfa2gff for MICE on 16 threads, while running the synteny block algorithms on one thread. Meanwhile we ran the entire minigraph-cactus (MG cactus) pipeline on 28 threads.

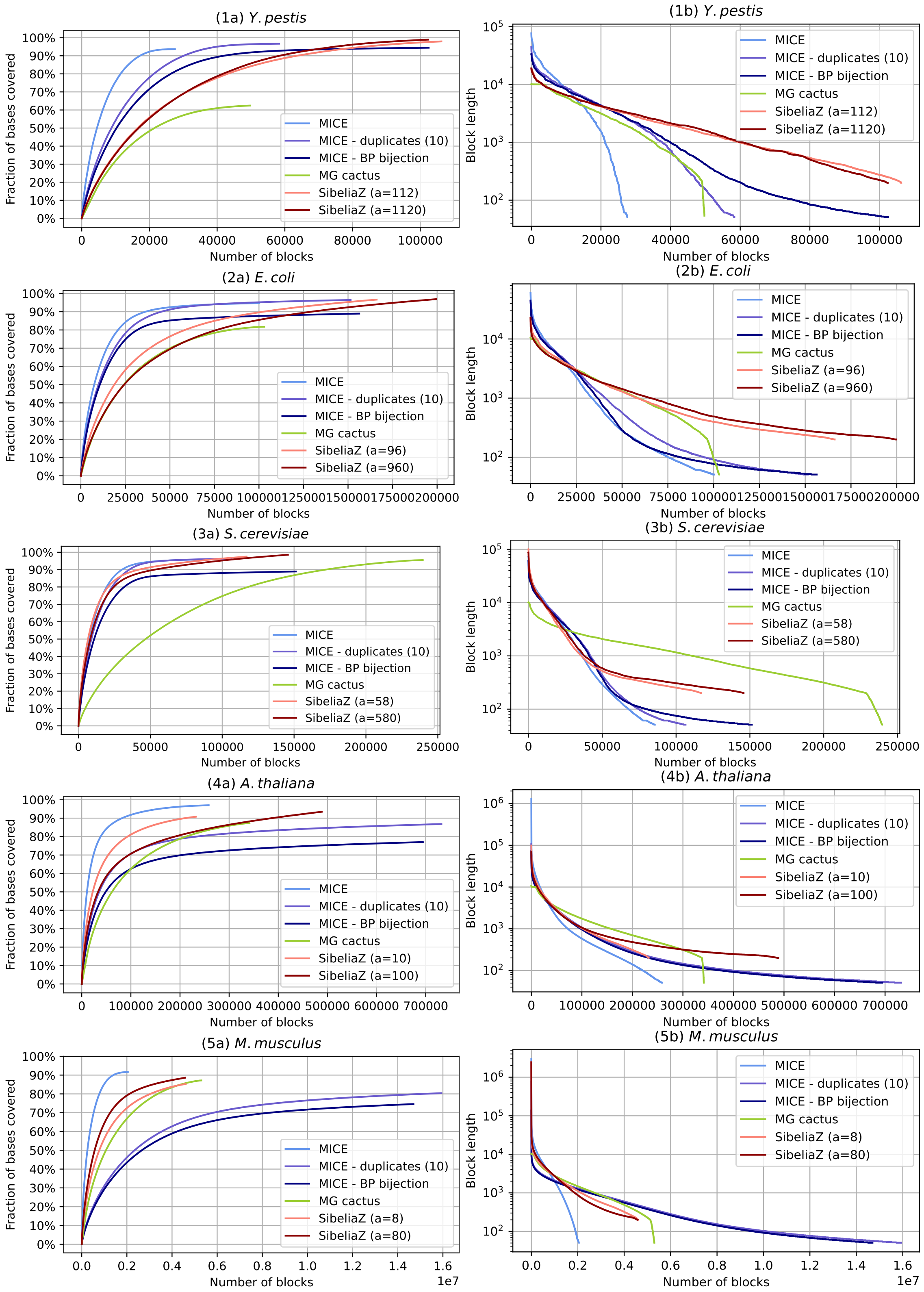


**Fig. 10.** Coverage of blocks (≥ 50 BP) and their size in base pairs in descending order of length generated by MICE, minigraph cactus and SibeliaZ on all 5 benchmark data sets.

**Table 2.** Encoding sizes as well as coverage across all genomes using block occurrences ≥ 50 BP in the five benchmark datasets.

| | *Y. pestis* | | *E. coli* | | *S. cerevisiae* | | *A. thaliana* | | *M. musculus* | |
|---|---|---|---|---|---|---|---|---|---|---|
| | enc. size | coverage | enc. size | coverage | enc. size | coverage | enc. size | coverage | enc. size | coverage |
| MICE | 27535 | 93.68% | 100182 | 94.73% | 85613 | 95.82% | 258296 | 97.02% | 2043614 | 91.67% |
| MICE - BP bijection | 102488 | 94.46% | 156282 | 88.97% | 151332 | 88.93% | 694185 | 77.02% | 14694541 | 74.58% |
| MICE - duplicates | 58312 | 96.68% | 151457 | 96.47% | 106418 | 96.36% | 731918 | 86.81% | 15932964 | 80.40% |
| MG Cactus | 49723 | 62.44% | 102950 | 81.73% | 239559 | 95.51% | 341340 | 87.38% | 5298901 | 87.16% |
| SibeliaZ - $a_{\text{low}}$ | 106243 | 97.96% | 166098 | 96.65% | 116994 | 97.51% | 232359 | 90.78% | 4603390 | 85.27% |
| SibeliaZ - $a_{\text{high}}$ | 102369 | 98.94% | 199547 | 96.90% | 145731 | 98.56% | 488621 | 93.46% | 4574842 | 88.58% |

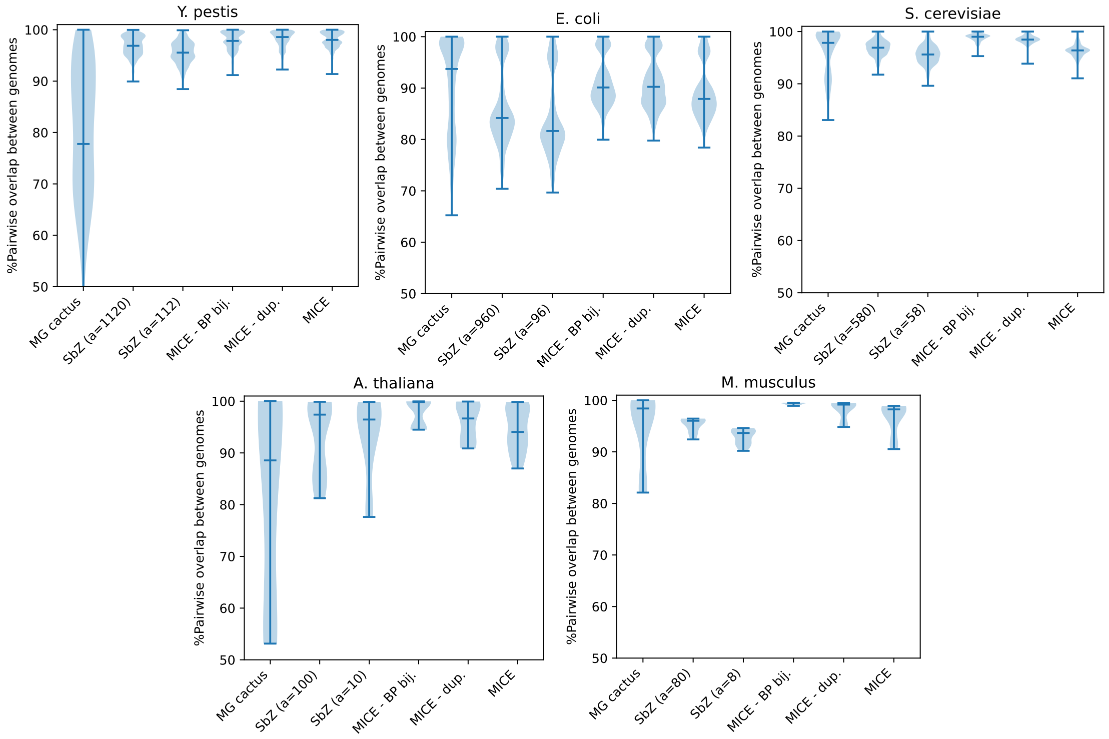


**Fig. 11.** Proportion of positions in shared blocks for pairs of genomes within those areas of the genome covered by blocks ≥ 50 BP.

### I Obscuring rearrangements

While MICE explicitly constructs a partition of the elements, the other tools do not. However, we can reconstruct which elements belong to the same block from the block identifiers in their output. Since these tools do not enforce a partition, some elements may be assigned to multiple block IDs.

We searched for two types of element breakpoints which are not recovered: (i) pairs of elements that form a breakpoint in at least one genome but appear together in at least one block with the same identifier, and (ii) breakpoints that occur entirely within a single block; the second type is a subset of the first. Examples of both types are shown in Appendix G, Fig. 7. In both cases, the breakpoint cannot be detected in downstream analyses when using synteny blocks alone.

Fig. 12 shows the percentage of breakpoints that are not obscured (blue), not recovered in the same block identifier (orange), and not recovered within the same phrase induced by a synteny block (red) for all methods. For MICE and MICE BP–bijection, no breakpoints are obscured by construction (Theorem 1). In MICE duplicates, where duplicates are considered for merging, some breakpoints are unrecoverable when both elements forming a breakpoint are merged with the same duplicate; this occurs in about 10–13% of cases. Although SibeliaZ uses a smaller $k$-mer size, both of its versions obscure 4–30% of breakpoints. Minigraph-Cactus obscures 8–16% of breakpoints and performs particularly poorly on *A. thaliana*, where 43% of breakpoints are not recovered.

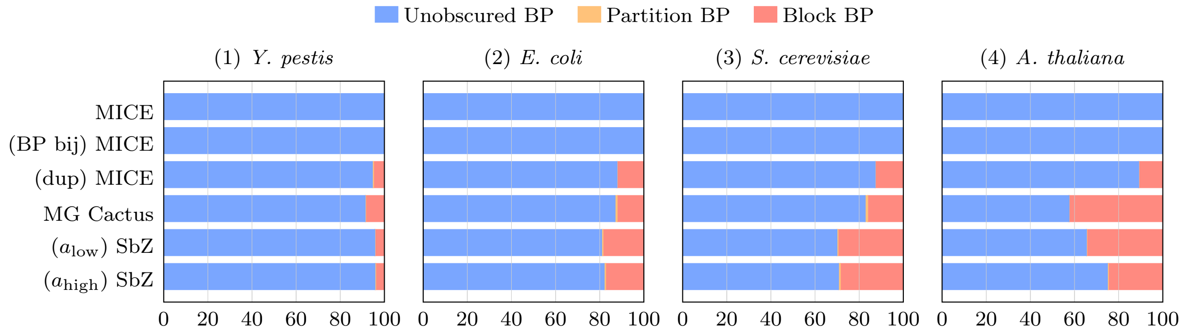


**Fig. 12.** Percentage of breakpoints that are not obscured (blue), not recovered by being placed in the same synteny block identifier (orange), or within the same phrase induced by a block (red), shown for all five methods (on the y-axis) and for four of the datasets. Table 3 lists the corresponding values.

**Table 3.** Comparison of breakpoints in a partition (Part) and breakpoints in a block (Block) across species for each method. The row "*Breakpoints*" represents the total number of breakpoints found from pairwise genome comparisons using 31-mers.

| | *Y. pestis* | | *E. coli* | | *S. cerevisiae* | | *A. thaliana* | |
|---|---|---|---|---|---|---|---|---|
| *Breakpoints* | 3945 | | 55867 | | 20957 | | 129338 | |
| | Part | Block | Part | Block | Part | Block | Part | Block |
| MICE | 0 | 0 | 0 | 0 | 0 | 0 | 0 | 0 |
| MICE - BP bijection | 0 | 0 | 0 | 0 | 0 | 0 | 0 | 0 |
| MICE - duplicates heuristic | 216 | 195 | 6815 | 6744 | 2679 | 2661 | 14129 | 14040 |
| MG-Cactus | 345 | 335 | 7305 | 6787 | 3628 | 3390 | 54967 | 54928 |
| SibeliaZ - $a_{low}$ | 172 | 167 | 10714 | 10362 | 6309 | 6203 | 44798 | 44494 |
| SibeliaZ - $a_{high}$ | 170 | 156 | 10073 | 9708 | 6158 | 5990 | 32493 | 31910 |

#### J A qualitative Example of a Locus in three *E. coli*

To give an impression about what the differences in marker contiguity and breakpoint preservation evaluated in Section 4 mean in practice, we repeated the experiment on *E. coli* with the first three genomes given in Table 1 and visualized a 40KB region that shows inverted, transposed and duplicated elements. The element matches are shown in Fig. 13.

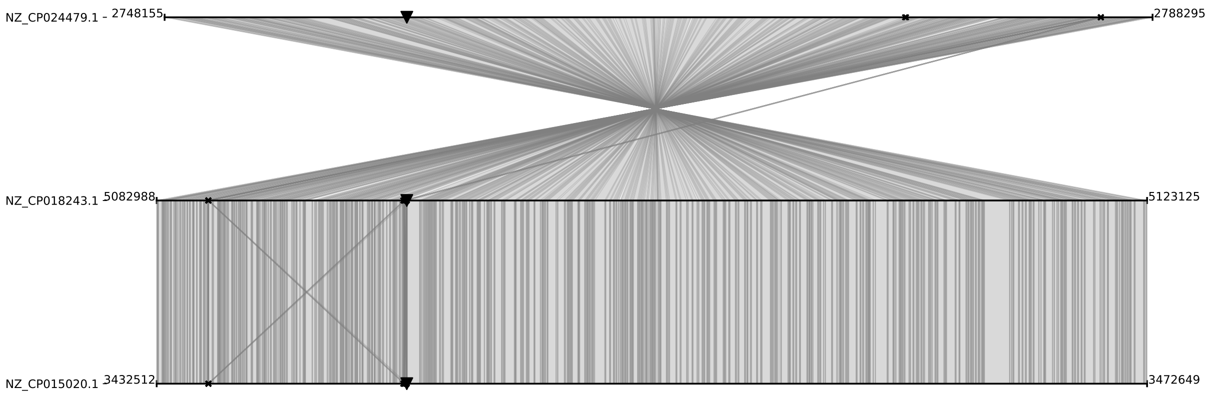


**Fig. 13.** 31-mer matches between three 40KB regions in three *E. coli* chromosomes. Crosses denote duplicate regions, triangles denote material that is found transposed outside of the shown window.

We show the synteny blocks determined by the different versions of MICE in Fig. 14. We see that all three versions of MICE recover the basic inversion and break blocks whenever a transposed element is observed. The main difference between the three versions of MICE is how duplicates are handled. They are ignored in the default mode (i), thus resulting in fewer and larger blocks. The duplicate mode (ii) considers the duplicate element anchors for their surrounding regions, i.e., it merges surrounding unique elements into the duplicates. The breakpoint bijection mode (iii) in contrast does not merge over the duplicates. Each duplicate is instead assigned its own marker, leaving the surrounding singular regions untouched.

Visualizing the blocks generated by SibeliaZ in Fig. 15, we observe that while with both abundance filters, SibeliaZ recovers the inversion, it needs more blocks to cover this region. Nonetheless, the block breaks themselves do not correspond to rearrangements observed in the $k$-mers. For example, in both cases, there are blocks that span transposed elements. In addition, some matches between duplicate regions are not retained.

We show the result for Minigraph-Cactus in Fig. 16. We observe that the blocks determined by Minigraph-Cactus still contain block breaks which do not seem to be directly induced by rearrangements and also do not correspond to block ends for SibeliaZ. However, none of the blocks shown here spans a transposed element. Nonetheless, the matches between certain copies of duplicates are similarly not preserved as for SibeliaZ.

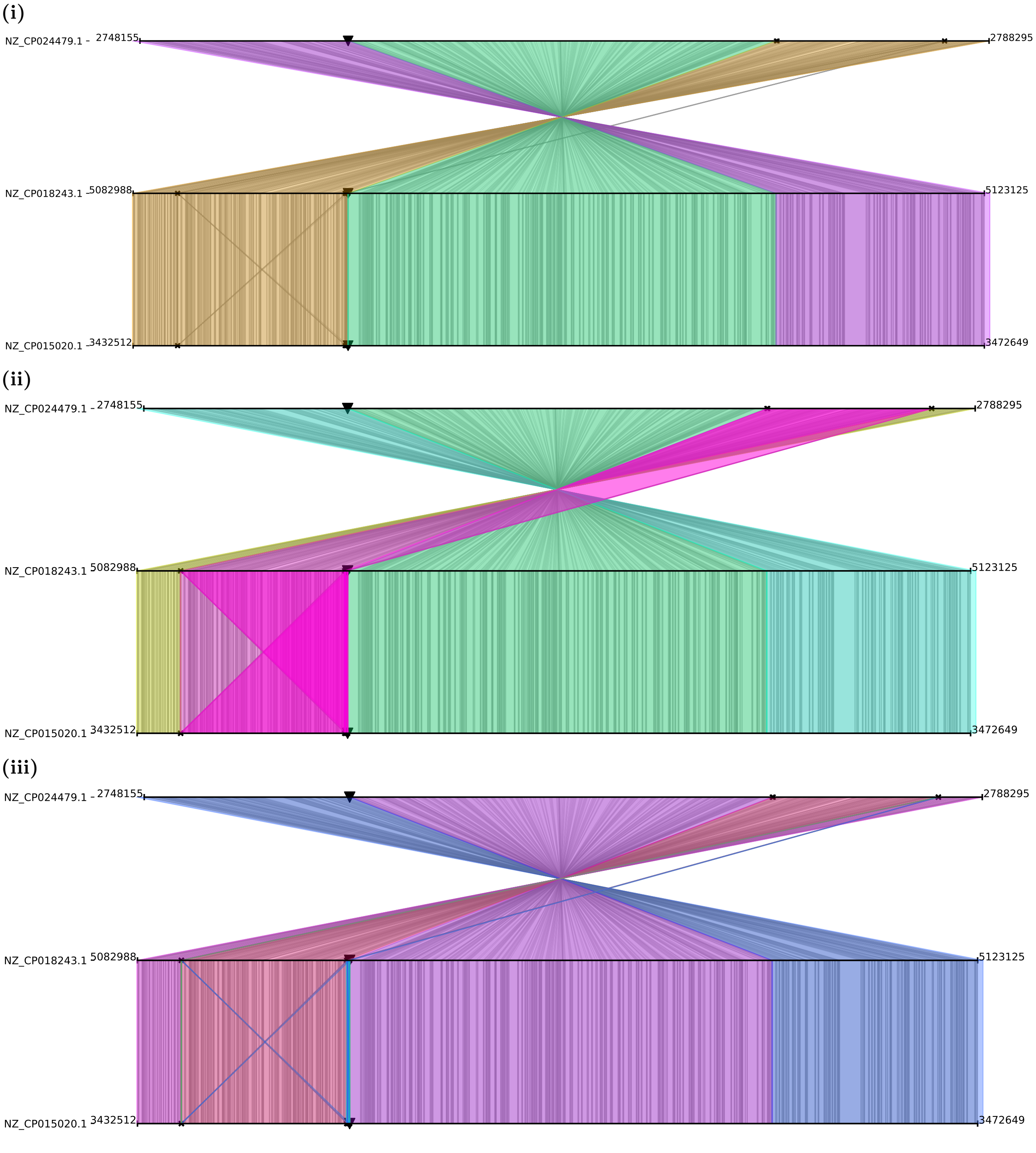


**Fig. 14.** Synteny blocks determined by MICE (i), MICE - duplicates (ii) and MICE - BP bijection (iii) overlayed on $k$-mer/unitig matches from Fig. 13.

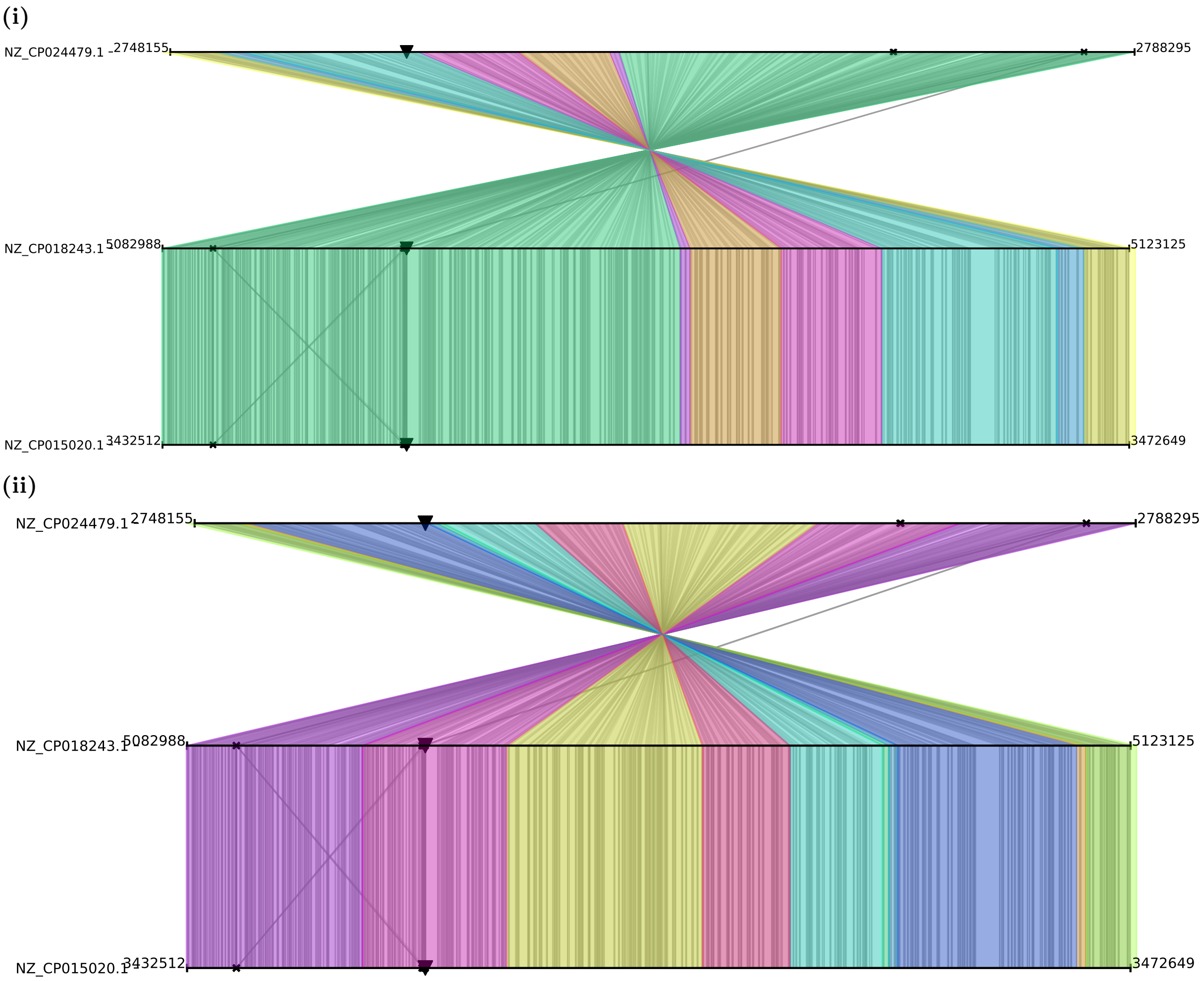


**Fig. 15.** Synteny blocks determined SibeliaZ with low abundance filter ($a_{\text{low}} = 6$) (i) and SibeliaZ with high abundance filter ($a_{\text{high}} = 60$) (ii) overlayed on $k$-mer/unitig matches from Fig. 13.

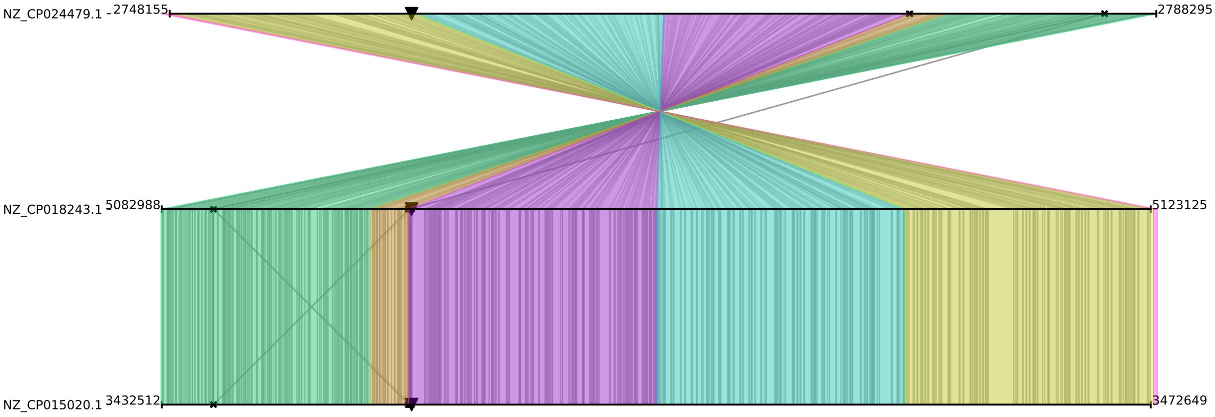


**Fig. 16.** Synteny blocks determined by Minigraph-Cactus overlayed on $k$-mer/unitig matches from Fig. 13.
